# Outbred *Drosophila* populations reveal diet-dependent genetic effects on development time

**DOI:** 10.64898/2026.06.05.730474

**Authors:** Yulin Bai, Sumaira Shabbir, Yuyan Chen, Fabio Morgante, Michael Ludwig, Soo-Young Park, Mohan Acharya, Yang I. Li, Sabir Ali, Abigail Trudnak, Mutheshree Rajesh, Martin Kreitman, Xuan Zhuang

**Author notes:** **Corresponding author:** Xuan Zhuang.

## Abstract

Metabolic effects of genetic variation often depend on diet, yet the loci underlying diet-dependent developmental responses remain incompletely defined. Here we combine multi-trait phenotyping of *Drosophila* Genetic Reference Panel lines with genome-wide mapping in newly developed *Drosophila* Recombinant Populations. High sugar causes genotype- and life-stage-dependent changes in metabolic and life-history traits, with development time emerging as a highly heritable, sugar-sensitive phenotype. Mapping in 16 outbred advanced intercross populations reveals distinct association landscapes under low- and high-sugar diets, with a concentrated low-sugar signal, a more distributed high-sugar pattern, and identified genotype-by-diet loci including *tap*, *Eip75B* and *Cerk*. Functional perturbation supports diet-dependent effects for several prioritized candidates. Allele-frequency analyses identify operationally defined thrifty-like variants associated with delayed development under high sugar and relatively earlier development under low sugar; these variants are enriched for cell-adhesion, neurodevelopmental, and morphogenetic processes. Together, these results establish an outbred *Drosophila* framework for dissecting how dietary sugar remodels the genetic architecture of development time.

## INTRODUCTION

Metabolic regulation is shaped by interactions between genetic variation and nutritional environment, yet much of the heritable basis of diet-responsive metabolic phenotypes remains unresolved despite the identification of numerous associated loci^1^. This problem is particularly relevant to type 2 diabetes, a disorder of glucose homeostasis in which genetic susceptibility and nutritional environment jointly shape disease risk. Because nutrient sensing and metabolic homeostasis influence growth, development, and survival, variants that compromise metabolic performance would often be expected to be consistently selected against^2^; nevertheless, natural populations retain substantial standing variation in metabolic performance. A plausible explanation is that many variants have conditional effects: their phenotypic consequences depend on diet, so costs in one nutritional regime can be offset by neutrality or benefits in another (genotype-by-environment interactions, G×E)^2–4^.

Specifically, the thrifty genotype hypothesis, originally proposed to explain why genetic susceptibility to diabetes might persist despite becoming detrimental in modern calorie-rich environments, proposes that alleles promoting efficient energy acquisition or storage may be favored under intermittent caloric scarcity but become disadvantageous under persistent caloric or sugar excess^5,6^. Although this hypothesis remains debated, it highlights a broader and experimentally testable idea: alleles can have environment-dependent consequences that are beneficial, neutral, or deleterious depending on nutritional context^7^. However, high-resolution maps that resolve how allelic effects change across controlled dietary conditions remain limited.

*Drosophila melanogaster* provides a powerful and experimentally scalable model for this purpose^8^, coupling conserved metabolic regulation with short generation time, large cohort sizes, and mature genetic tools under precisely defined diets. Importantly, these conserved nutrient-responsive pathways and metabolic programs^9–12^, including insulin/ insulin-like growth factor (IGF) and target of rapamycin (TOR) signaling as well as core carbohydrate and lipid metabolism, regulate not only metabolic homeostasis but also growth and developmental progression. As a result, high-sugar diet exposure in *Drosophila* induces not only metabolic abnormalities relevant to diabetic physiology such as hyperglycemia, lipid accumulation, and insulin resistance, but also organismal consequences including developmental delay and reduced survival^2,9,13–15^.

To connect diet-responsive metabolic and life-history variation with its underlying genetic basis, we adopted a complementary two-stage strategy in *Drosophila*. We first used inbred lines from the *Drosophila* Genetic Reference Panel (DGRP)^16^, which enable repeated measurements of the same genotype across dietary conditions, to profile a panel of metabolic and life-history traits under a low-sugar diet (LSD) and a high-sugar diet (HSD), and to characterize their genetic architecture, heritability, and correlation structure across environments. This phenotypic survey highlighted development time as a particularly informative diet-responsive trait. We then developed *Drosophila* Recombinant Populations (DRPs), an outbred, advanced-intercross mapping resource composed of 16 large, highly recombined populations with repeated founder representation across populations (**Figure 1**). Because these populations were derived from defined founder sets and maintained through extended intercrossing, the design combines increased recombination with informative founder-allele frequencies and cross-population reproducibility, and is well suited for extreme-phenotype sequencing^17^. Using this framework together with genome sequencing of more than 6,000 flies, we dissect the genetic basis of development time under low-and high-sugar diets. This analysis reveals that development time has a diet-dependent genetic architecture, with largely distinct loci under LSD and HSD as well as loci showing gene-by-diet effects. Functional analyses further supported candidate regulators of diet-dependent variation in development time.

**Figure 1.**
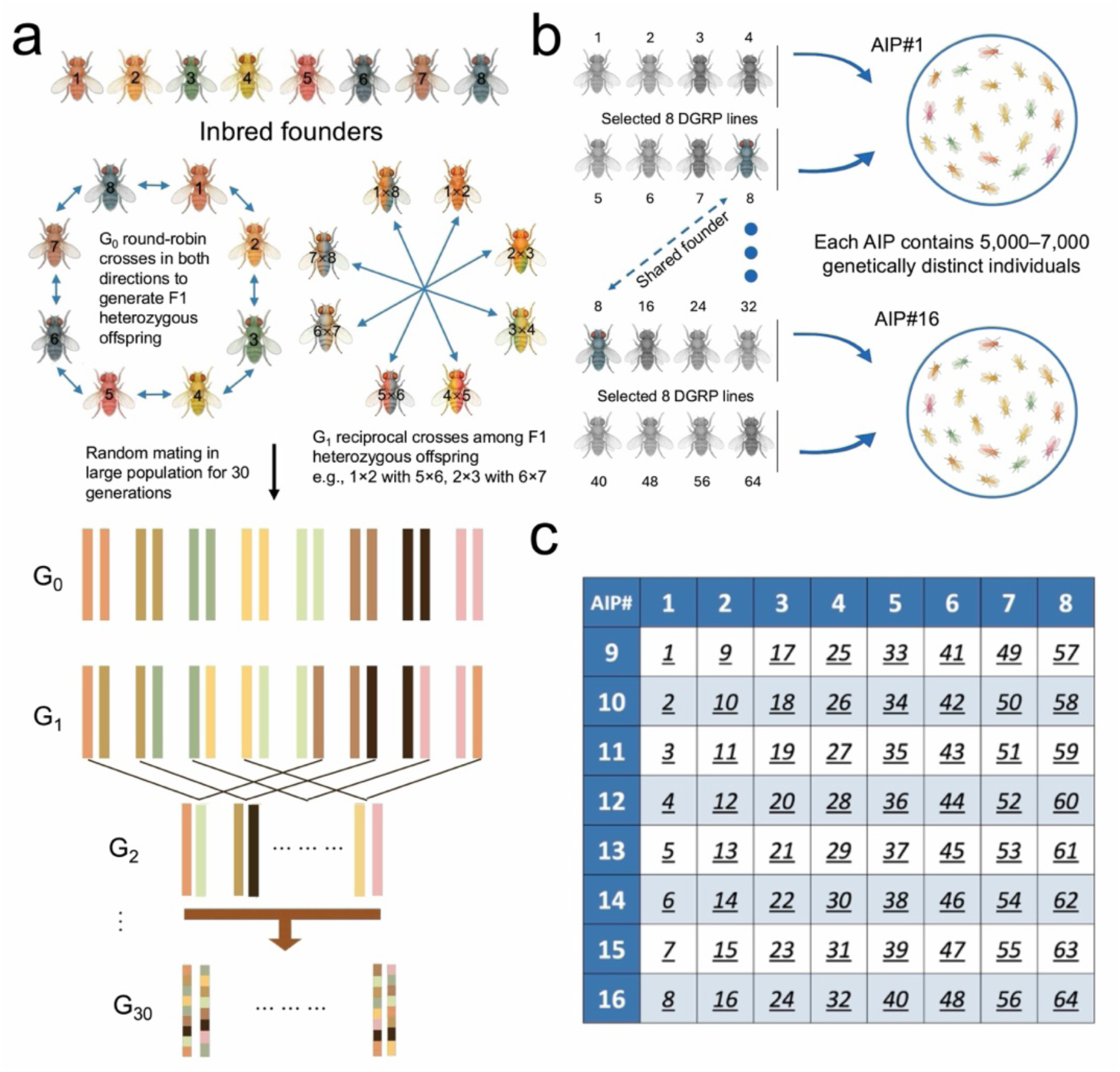
Construction of the *Drosophila* Recombinant Populations (DRPs). **a**, Schematic of DRP construction from *Drosophila* Genetic Reference Panel (DGRP) founders. Sixty-four inbred DGRP lines were used to establish 16 outbred advanced intercross populations (AIPs), each initiated from eight founder lines. Coloured bars represent founder-derived chromosomes across generations. Founder chromosomes were mixed through controlled crosses and subsequent random mating, and recombination was allowed to accumulate for at least 30 generations before mapping. **b**, Example founder sets for AIP1 and AIP16. Each AIP was established by pooling descendants from eight designated DGRP lines and maintained as an outbred population of approximately 5,000–7,000 genetically distinct individuals. The dashed arrow indicates a founder line shared between the two example AIPs. **c**, Founder-allocation matrix for the 64 DGRP lines and 16 AIPs. White bold numbers indicate AIPs, and underlined italic numbers indicate DGRP founder lines. The eight founders in each column or row form the AIP indicated at the top or left, respectively. Each founder appears in two AIPs, creating founder overlap across the DRP resource for cross-population comparison.

## RESULTS

### DGRP reveals stage- and genotype-dependent sugar responses

To investigate how genetic background influences phenotypic responses to dietary sugar, we measured seven metabolic and life-history trait classes, comprising 11 stage-specific phenotypes, in larvae and adults from 32 DGRP lines included among the DRP founders (**Supplementary Table 1**) and reared under HSD and LSD dietary conditions (**Supplementary Data 1**). Traits quantified included whole-body glucose, glycogen, and triglyceride levels, as well as body weight, development time, larval survival, and adult longevity (**Figure. 2a**).

**Figure 2.**
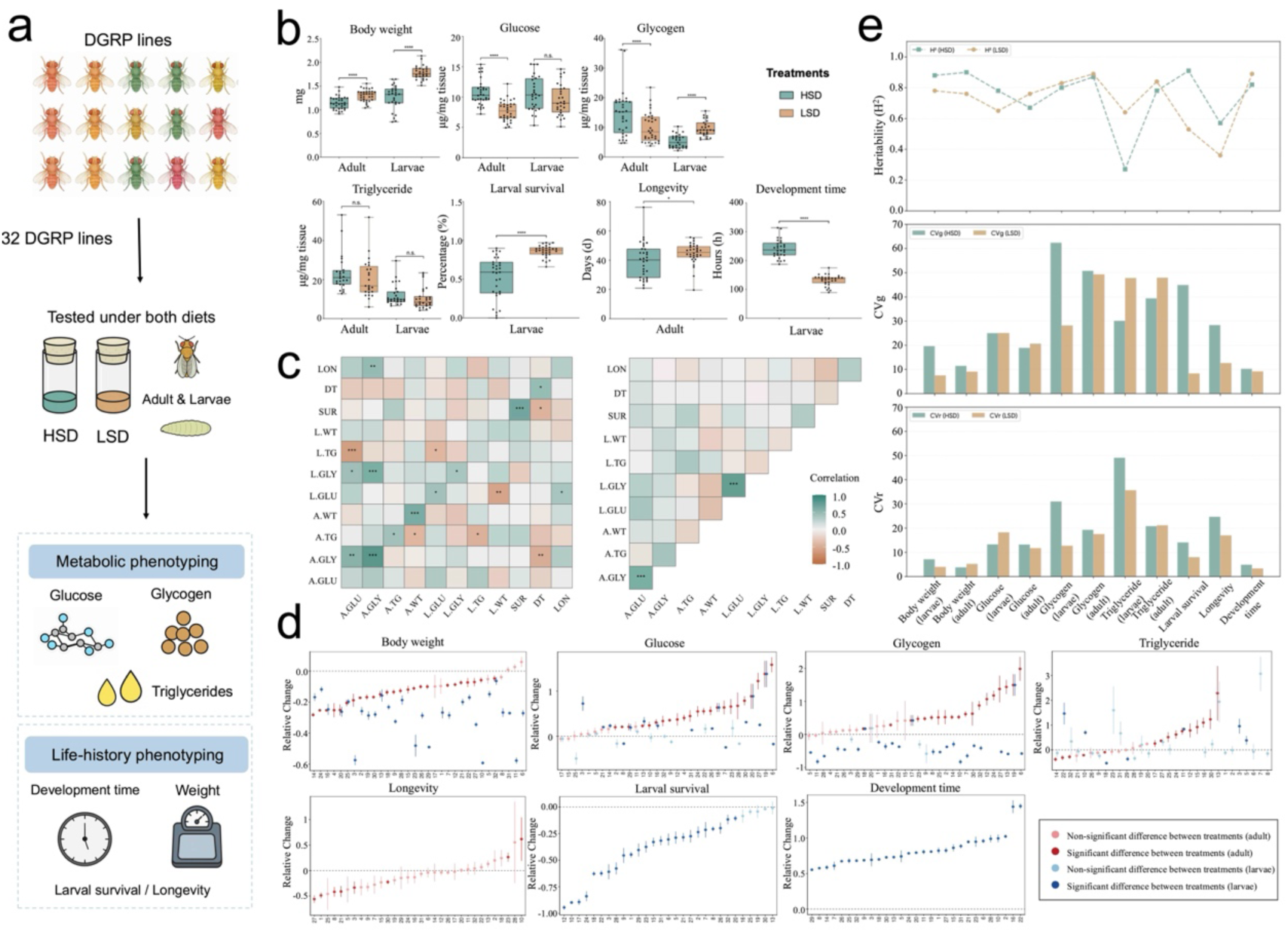
Diet-dependent phenotypic variation across DGRP lines. **a**, Experimental design for phenotyping 32 DGRP lines included among the DRP founders under high-sugar diet (HSD) and low-sugar diet (LSD). Metabolic traits included whole-body glucose, glycogen, and triglyceride levels; life-history traits included body weight, development time, larval survival, and adult longevity, measured in larvae and/or adults as indicated. **b**, Distributions of metabolic and life-history traits under HSD and LSD. Box plots show variation across DGRP line means; points indicate individual line means. Significance comparisons between diets were performed using ANOVA. ns, not significant; \**P* < 0.05; \*\**P* < 0.01; \*\*\**P* < 0.001; \*\*\*\**P* < 0.0001. **c**, Spearman correlation matrices for diet responses and diet-specific phenotypes. In the left heatmap, the upper triangle shows correlations among traits under LSD, the lower triangle shows correlations among traits under HSD, and the diagonal shows cross-diet correlations for the same trait. The right heatmap shows correlations among relative diet responses, calculated as (HSD − LSD)/LSD. Positive correlations are shown in green and negative correlations in brown. Significant correlations are indicated by asterisks. A, adult; L, larval; WT, body weight; GLU, glucose; GLY, glycogen; TG, triglyceride; SUR, survival; LON, longevity; DT, development time. **d**, Relative phenotypic changes between HSD and LSD across DGRP lines. Points show line means, and error bars indicate the standard error of the relative change. Adult traits are shown in red and larval traits in blue; darker points indicate significant diet differences. For traits measured in both stages, lines are ordered by the adult response. **e**, Broad-sense heritability and scaled variance measured under HSD and LSD. The top panel shows broad-sense heritability (H²), calculated as the proportion of between-line variance relative to total phenotypic variance. The middle and bottom panels show genotypic coefficient of variation (CV_G_) and residual coefficient of variation (CV_R_), respectively, after scaling variance components by trait means

In the adult experiments, all lines survived the 5-day dietary treatment under both HSD and LSD conditions. In the larval experiments, developmental failure itself emerged as an informative phenotype: five lines (#4, #12, #15, #17, and #24) (**Supplementary Table 1**) failed to complete development on HSD, and line 15 also failed to develop on LSD. Rather than representing missing observations alone, these lines highlight extreme genotype-specific sensitivity to dietary conditions.

On average, flies reared on HSD exhibited elevated glucose levels, reduced body weight, delayed development, and shortened survival and lifespan relative to those reared on LSD (**Figure 2b**), consistent with prior *Drosophila* models of diet-induced metabolic disruption^9,15,18^. To further characterize these responses across genotypes, we calculated the relative change between diets, (HSD−LSD)/LSD, to summarize each line’s sugar response and assess both the direction and magnitude of each line’s phenotypic response (**Figure 2d, Supplementary Data 1**). Among the 11 traits examined, only four—larval body weight, larval glycogen, development time, and larval survival rate—showed a consistent directional response across all genotypes on HSD relative to LSD. In contrast, the remaining seven traits showed bidirectional, genotype-dependent responses. Triglyceride levels in both larvae and adults displayed the clearest bidirectional responses across diets. Except for larval triglyceride levels, all traits showed significant diet effects (**Figure 2b**). All larval and adult traits also exhibited significant line effects within each diet, and diet × line interactions were significant for all traits (**Supplementary Table 3**), demonstrating that genetic background strongly modulates the phenotypic response to dietary sugar concentration. Together, these results indicate that HSD induces broad phenotypic shifts whose magnitude and, for several traits, even direction depend strongly on genotype, thereby motivating the subsequent diet-dependent and G×E mapping analyses.

### Trait correlations differ across diets and life stages

We first examined correlations among trait values within and across diets (**Figure 2c, left**). Many same traits were positively correlated between HSD and LSD, indicating partial preservation of genotype-dependent trait differences across dietary environments. Diet-specific correlations revealed distinct patterns. Under HSD, larval glucose was inversely correlated with larval body weight, and adult triglyceride was inversely correlated with adult body weight, suggesting that sugar- or lipid-associated metabolic load is linked to reduced body mass under high-sugar stress. Development time was also negatively correlated with larval survival and adult glycogen under HSD, linking slower development with reduced larval performance and lower carbohydrate storage. Under LSD, carbohydrate traits were more tightly coupled: adult glucose was positively correlated with adult glycogen and larval glycogen, whereas larval triglyceride was negatively correlated with glucose traits. Together, these correlations suggest that trait relationships shift with diet, with HSD emphasizing links between metabolic stress and life-history performance, and LSD showing stronger coordination among carbohydrate-storage traits.

We next examined correlations in relative diet response, calculated as (HSD−LSD)/LSD (**Figure 2c, right**). These response correlations were fewer and largely stage-specific. The clearest pattern was the positive association between glucose and glycogen responses within each life stage: lines with larger diet-induced changes in glucose also tended to show larger changes in glycogen in both adults and larvae. For traits measured in both larvae and adults, the magnitude of the diet response was not strongly correlated across life stages, indicating that sugar-responsive trait architecture is organized mainly within developmental stage rather than shared uniformly across larvae and adults.

### High-sugar stress exposes cryptic variation

The diet responses of the examined DGRP lines spanned a continuous range rather than discrete phenotypic classes (**Figure 2d**), indicating quantitative genetic variation across genotypes and supporting the suitability of these traits for genetic mapping.

Estimates of broad-sense heritability (H²) ranged from 0.31 to 0.90 across traits (**Figure 2e, Supplementary Table 4**), with most values exceeding 0.5. In general, H² was higher under HSD than under LSD, indicating that high-sugar stress increases the relative contribution of genetic differences to phenotypic variation. Notably, larval survival and development time were among the most heritable traits, making them particularly informative for genetic mapping. Across most traits, between-line genetic variance (Vg), within-line residual variance (Vr), and total phenotypic variance were all greater under HSD than under LSD, with triglyceride level as the main exception. After scaling variance components by trait means, genotypic coefficients of variation (CVg) ranged from 7.71% to 63.60%, whereas residual coefficients of variation (CVr) ranged from 3.24% to 55.74% and were generally lower than CVg (**Figure 2e, Supplementary Table 4**). The broader phenotypic spread under HSD is consistent with diet-dependent decanalization^19,20^, in which high-sugar stress reduces developmental buffering and allows genetic or individual-level variation that is constrained under LSD to become expressed. Consistent with the idea that the balance between homeostasis and plasticity can be modified by environmental conditions^21^, HSD appears to shift development away from a more canalized state, producing greater dispersion of phenotypic outcomes. These results show that the phenotypic effects of dietary sugar are strongly background dependent and support a model in which HSD enhances trait variance through genotype-specific responses, G×E interactions, and diet-dependent decanalization.

Among these traits, development time emerged as the most informative for subsequent genetic analysis. It showed one of the highest broad-sense heritability estimates (**Figure 2e, Supplementary Table 4**) and was the only measured trait with non-overlapping HSD and LSD distributions in the box plots (**Figure 2b**). Together with its expanded variance under HSD, these features indicates that development time is both highly diet-sensitive and strongly genetically influenced, making it the most suitable phenotype for subsequent genetic mapping in the DRPs.

### Diet-specific meta-GWAS identifies development time loci

To move from broad phenotypic characterization in the DGRP to higher-powered genetic dissection of development time, we established the DRPs as 16 outbred advanced intercross populations (AIPs). This design was intended to leverage large population size, accumulated recombination, and repeated founder representation across populations to improve detection of rare founder variants in the DGRP, while also increasing mapping resolution and enabling reproducibility to be assessed across populations. Within each population and diet, we used eclosion time to define phenotypic extremes and collected the earliest- and latest-eclosing flies for mapping. We then applied meta-GWAS framework integrating founder-haplotype inference and cross-population reproducibility filtering to identify loci affecting development time under LSD and HSD (**Figure 3a; see methods for details**).

**Figure 3.**
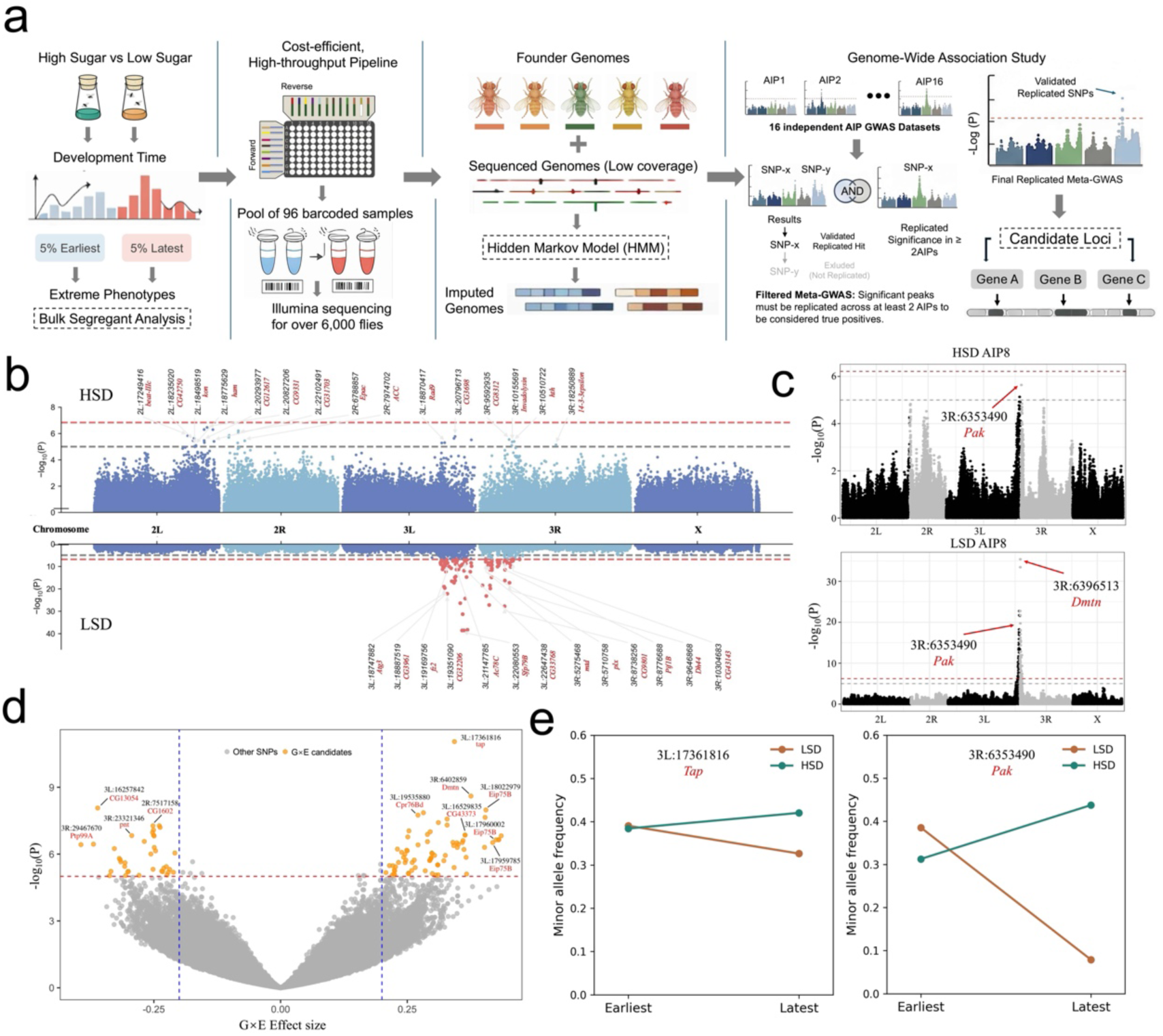
DRPs-based extreme-phenotype mapping identifies diet-dependent loci associated with development time. **a**, Overview of the DRP mapping strategy. For each AIP and diet condition, flies from the earliest and latest 5% of eclosion time were selected as phenotypic extremes. In total, 6,144 individually barcoded flies were pooled into 96-sample sequencing libraries. Founder haplotypes were inferred from low-coverage sequence data and founder genomes using a hidden Markov model (HMM), followed by AIP-specific GWAS, meta-analysis across AIPs, and reproducibility filtering to retain replicated candidate loci. **b**, Manhattan plots of diet-specific meta-GWAS for development time under HSD and LSD. The x-axis shows genomic position by chromosome arm, and the y-axis shows −log₁₀(*P*). Dashed lines indicate nominal (gray) and Bonferroni-adjusted (red) significance thresholds. Representative loci are labelled by genomic positions and related-candidate genes. **c**, Representative AIP-specific GWAS profiles for AIP8 under HSD and LSD. Red arrows indicate peak SNPs near *Pak* and *Dmtn*. **d**, Genome-wide genotype-by-diet interaction analysis. The volcano plot shows G×E effect estimates and −log₁₀(P). Orange points indicate SNPs passing the indicated significance and effect-size criteria; grey points indicate all other SNPs. The red dashed line indicates the nominal significance threshold (*P* < 1×10^−5^), and blue dashed lines indicate effect-size cutoffs (∣G×E *effect*∣ > 0.2). Representative G×E loci are labelled. **e**. Diet-dependent allele-frequency shifts between early- and late-eclosing flies. Minor allele frequencies are shown for representative SNPs near *Tap* and *Pak*.

Diet-specific meta-GWAS revealed markedly different association patterns under LSD and HSD, with most significant loci showing little overlap between the two dietary conditions (**Figure 3b, Supplementary Figure 3**). Under LSD, 104 SNPs exceeded the Bonferroni-adjusted significance threshold (*P* < 1.47 × 10⁻⁷), and 174 SNPs exceeded the nominal significance threshold (*P* < 1 × 10⁻⁵)^22^ (**Supplementary Data 2**). These associations were dominated by a strong and highly concentrated signal spanning the centromeric region of chromosome 3, indicating that much of the detectable signal under LSD is localized to a restricted genomic interval. In contrast, under HSD, no SNP exceeded the Bonferroni-adjusted significance threshold (*P* < 1.45 × 10⁻⁷), and only 27 SNPs exceeded the nominal significance threshold (**Supplementary Data 2**). Instead, the HSD signal was more dispersed, with significant or near-significant SNPs distributed across multiple regions on chromosomes 2 and 3. Unlike the prominent chromosome 3 cluster observed under LSD, the HSD pattern lacked a single dominant block and instead consisted of several smaller peaks, consistent with a more distributed genetic architecture under sugar stress. Polygenic score (PGS) analyses further supported this interpretation as prediction improved when broader sets of SNPs were included, indicating that development time is influenced by many variants of modest effect under both LSD and HSD (**Supplementary Figure 5**). Thus, although some loci rose above the genome-wide background, development time appears to reflect both detectable association peaks and a broader polygenic component.

The lead association under LSD mapped near *Sfp79B*, within the major chromosome 3 centromeric peak, and the top SNP reached *P* = 2.56 × 10⁻³⁹. Additional annotated loci within this broad interval included *Atg3*, *CG9801*, *fz2*, *CG32206*, *Ac78C*, *CG33768*, *mtd*, *plx*, *Pif1B*, *D144*, and *CG43143*. Several genes among them have functions relevant to growth and developmental regulation, including *Atg3*, a core autophagy gene; *Ac78C*, an adenylyl cyclase involved in cAMP signaling; *plx*, implicated in developmental patterning; and mtd, linked to immune and stress-responsive processes.

In contrast, under HSD, the lead association mapped near *CG9331*, with the top SNP reaching *P* = 3.46 × 10⁻⁷. The strongest HSD signals were distributed across several distinct genomic regions rather than concentrated within a single interval. Annotated loci included *beat-IIc*, *CG42750*, *ham*, *CG12617*, *CG31703*, *Epac*, *ACC*, *Rady*, *CG3698*, *CG8312*, *Invadolysin*, *hth*, and *14-3-3epsilon*. Several of these genes have functions consistent with environmentally sensitive developmental responses, including *Epac*, a cAMP-responsive signaling factor; *ACC*, which encodes acetyl-CoA carboxylase, a key enzyme in fatty acid biosynthesis; *Invadolysin*, a metalloprotease implicated in cell migration and chromosome organization; *hth*, a developmental regulator; and *14-3-3epsilon*, a conserved modulator of multiple signaling pathways. Together, these results indicate that development time under HSD is influenced by a broader and more distributed set of loci involving signaling, metabolism, and developmental regulation.

In addition to the across-population meta-analysis, population-specific scans revealed heterogeneity in the loci detected within individual DRP populations (**Figure 3c, Supplementary Figure 4**), consistent with the expectation that some genetic effects are background dependent. These AIP-specific associations illustrate the value of analyzing individual DRP populations separately, as such signals may be diluted or masked when effects are combined across independently derived populations.

Population-specific scans also revealed diet-dependent changes in allelic effects. For example, in AIP 8, the SNP (3R;6353490) near pak was associated with development time under both diets but showed opposite effect directions, with a positive effect under HSD (effect size = 0.24, *P* = 2.30 × 10⁻⁶) and a negative effect under LSD (effect size = −0.40, *P* = 1.85 × 10⁻²⁰). Together, these results show that the DRP design captures both background-dependent signals across populations and diet-dependent reversals of genetic effects within populations.

### Genotype-by-diet mapping identifies interaction loci

We next tested genotype-by-diet interaction effects by combining the LSD and HSD datasets in a meta-GWAS framework (**Figure 3d**). This analysis identified 122 SNPs exceeding the nominal significance threshold for G×E effects on development time, of which 21 remained significant after Bonferroni adjustment (**Supplementary Data 2**). These G×E-associated variants were distributed on both sides of the interaction-effect axis, indicating that allelic effects could reverse direction between diets, beyond simple changes in magnitude. Comparison with the separate diet-specific scans showed limited overlap: only one G×E-associated SNP overlapped with the HSD scan, whereas 13 overlapped with the LSD scan (**Supplementary Figure 3**). Thus, most interaction signals were not simply rediscovered in the separate diet-specific analyses, indicating that their effects are better captured by explicit modeling of dietary contrast.

The strongest positive G×E signal was detected near tap, while additional high-ranking loci included *Eip75B*, *Dmtn*, *Cpr76Bd*, *CG43373*, *pak*, *Cerk*, *CG1602*, *pnt*, *Ptp99A*, and *CG13054*. Although these candidates do not converge on a single canonical pathway, several are connected by functions in developmental regulation, morphogenesis, endocrine regulation, and signal transduction, suggesting that diet-responsive variation in development time may arise from coordinated effects on growth regulation and tissue organization. Among these candidates, *Eip75B* is particularly notable because it encodes an ecdysone-induced nuclear receptor involved in the hormonal regulation of developmental transitions. Its recovery in the G×E scan is consistent with the role of endocrine signaling in linking nutritional state to developmental progression, as well as with prior evidence implicating *Eip75B* in nutrient-responsive metabolic programming, including sugar-stress or lipogenic outputs^23^. Other high-ranking candidates also point to related developmental and signaling processes. For example, *tap* and *pak* are linked to neuronal development, axon growth, cytoskeletal remodeling, and morphogenesis, whereas *Cerk* and *pnt* implicate sphingolipid signaling and receptor tyrosine kinase/mitogen-activated protein kinase (RTK/MAPK)-associated regulation. Additional candidates such as *CG13054*, *CG1602*, and *CG43373* remain less well characterized, but their strong interaction signals highlight potentially underappreciated contributors to sugar-responsive developmental variation.

We next examined allele-frequency shifts at representative loci to visualize these interaction effects (**Figure 3e**). At the top G×E locus near *tap*, the minor allele decreased in frequency from the earliest to the latest developers under LSD but increased under HSD, producing a clear crossover pattern. A similar diet-dependent pattern was observed near *pak*. These allele-frequency shifts provide a direct illustration of G×E.

### Functional validation reveals context-dependent gene effects

To functionally validate candidate loci identified by the diet-specific GWAS and combined G×E analysis, we tested prioritized genes using available *Drosophila* perturbation lines (**Figure 4a**), including RNAi knockdown (KD), overexpression (OE), and independent loss-of-function (LOF) alleles. Based on mapping support, annotation, and stock availability, we selected seven G×E-prioritized genes (*Eip75B*, *Cerk*, *pak*, *tap*, *CG43373*, *Dmtn*, and *Cpr76Bd*), one HSD-prioritized candidate (*CG9331*), and ten LSD-prioritized candidates (*CG33768*, *CG9801*, *CG3961*, *CG32206*, *plx*, *Pif1B*, *mtd*, *Sfp79B*, *fz2*, and *Dh44*). Overall, the validation experiments provided evidence of developmental roles for 16 of the 18 selected candidates: four knockdown candidates produced perturbation-specific pre-adult lethality, and 12 of the 14 quantitatively assayed candidates significantly altered development time. Among the five G×E-prioritized candidates evaluated under both diets, four showed diet-dependent effects; four quantitatively assayed genes also showed sex-specific effects, with two displaying both diet- and sex-dependent patterns (**Supplementary Figure 6, Supplementary Figure 7, Supplementary Table 5**).

**Figure 4.**
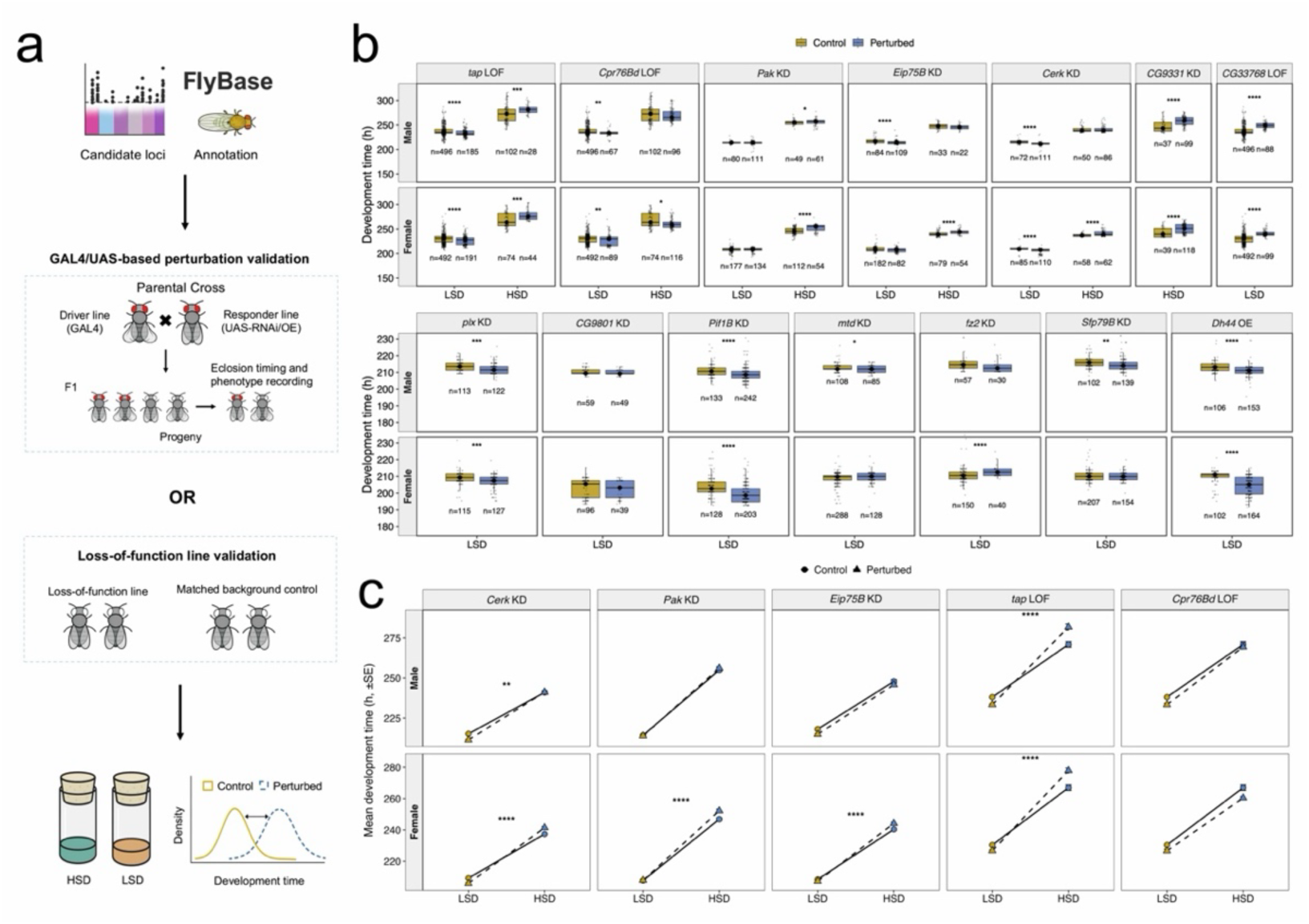
Functional validation of candidate genes. **a**, Schematic of the validation workflow. Candidate loci identified from GWAS were annotated using FlyBase and tested using either UAS/GAL4-based perturbation assays or independent loss-of-function lines. For UAS-based assays, GAL4 driver lines were crossed to responder lines carrying UAS-RNAi knockdown (KD) or UAS-overexpression (OE) constructs, and F1 progeny were assigned to perturbed or sibling-control classes. For loss-of-function (LOF) assays, candidate lines were compared with matched genetic-background control lines. Egg-to-adult development time was recorded under HSD and/or LSD, depending on the candidate class. **b**, Development-time distributions for candidate-gene perturbation assays. Box plots show egg-to-adult development time for control and perturbed genotypes, separated by candidate gene, sex and diet. Yellow boxes indicate controls, and blue boxes indicate perturbed genotypes and grey points indicate individual flies. Sample sizes below each box indicate the number of flies scored. Asterisks indicate two-sided Wilcoxon rank-sum tests comparing perturbed and matched-control individuals within each sex and diet. **c**, Diet-dependent perturbation effects for selected G×E-prioritized candidates tested under both diets. Points show mean egg-to-adult development time, and error bars indicate standard error (s.e.). Control and perturbed genotypes are plotted separately for each candidate gene and sex, with lines connecting the same genotype group across diets. Asterisks indicate the perturbation status × diet interaction from linear models fitted separately for each candidate gene and sex. \**P* < 0.05; \*\**P* < 0.01; \*\*\**P* < 0.001; \*\*\*\**P* < 0.0001

RNAi knockdown of four candidate genes—*CG43373*, *Dmtn*, *CG3961*, and *CG32206*—resulted in complete pre-adult lethality, with no recovered adults carrying the intended perturbation under either diet, whereas the corresponding sibling control genotypes remained viable (**Supplementary Data 3**). These results indicate that the lethality was specifically associated with knockdown of these genes in the experimental genotypes. These results suggest that the affected genes are required for early viability or development before adult eclosion. This is consistent with the known involvement of *Dmtn* in cytoskeletal organization, while the less characterized genes *CG43373*, *CG3961*, and *CG32206* may also have essential, diet-independent roles in early viability. For the remaining 14 candidates, egg-to-adult development time was measured relative to matched controls: sibling F1 controls for KD and OE assays, and genetic-background-matched controls for LOF assays. Diet-specific candidates were assayed under the dietary condition in which they were originally identified, whereas G×E-prioritized candidates were evaluated under both LSD and HSD to test diet-dependent effects directly (**Supplementary Data 4**).

Across sexes and diets, candidate-gene perturbations showed distinct patterns of effects on egg-to-adult development time (**Figure 4b, Supplementary Figure 7, Supplementary Table 5**). Several candidates, including *plx*, *CG9331*, *mtd*, *fz2*, *Cpr76Bd*, and *CG33768*, showed significant perturbation effects without detectable diet- or sex-specific modification, indicating broadly concordant perturbation effects across the tested groups. In contrast, other candidates showed context-dependent effects that were restricted to a specific sex, modified by diet, or dependent on both sex and diet. Among the G×E-prioritized candidates, *Cerk* RNAi and *Eip75B* RNAi showed combined diet- and sex-specific effects. *Cerk* RNAi shortened development time under LSD but increased development time under HSD, with the sex-specific effect detected under HSD. *Eip75B* RNAi showed a diet-specific effect in females and a sex-specific effect under HSD, consistent with a sex-by-diet-dependent perturbation pattern. *Pak* RNAi and tap LOF showed diet-specific effects without detectable sex-specific effects; *Pak* showed a diet-specific effect in females, with tap showing diet-specific effects in both sexes. In contrast, *Cpr76Bd* LOF accelerated development overall, with no detectable diet- or sex-specific modification. Among candidates assessed under LSD, *Pif1B* RNAi and *Dh44* overexpression showed sex-specific effects, whereas *CG9801* RNAi and *Sfp79B* RNAi did not significantly alter development time. Together, these results indicate that mapped candidates influence development time through both broad developmental effects and context-dependent responses shaped by diet and sex (**Figure 4c, Supplementary Table 5**).

### Directional diet-opposed shifts identify thrifty-like SNPs

The thrifty genotype hypothesis proposes that alleles advantageous under caloric scarcity may become costly under caloric excess. By analogy with this framework, we defined “thrifty-like” variants in our study as alleles enriched in late-eclosing flies under HSD but depleted in late-eclosing flies under LSD, corresponding to variants associated with delayed development under HSD but relatively earlier development under LSD.

To identify such variants, we compared allele-frequency shifts between early- and late-eclosing flies within each diet and screened for SNPs showing this directional diet-opposed pattern (**Supplementary Figure 8**). This contrast-based analysis highlights SNPs with directly interpretable cross-diet reversals at the allele level, complementing the model-based G×E analysis. Using this framework, we identified 2,674 candidate SNPs showing contrasting allele-frequency changes between diets (**Figure 5a**). Comparison with a randomized null distribution then highlighted a subset of 1082 SNPs with strong support for this directional diet-opposed pattern, which we retained as robust thrifty-like variants (**Figure 5b, Supplementary Data 5**).

**Figure 5.**
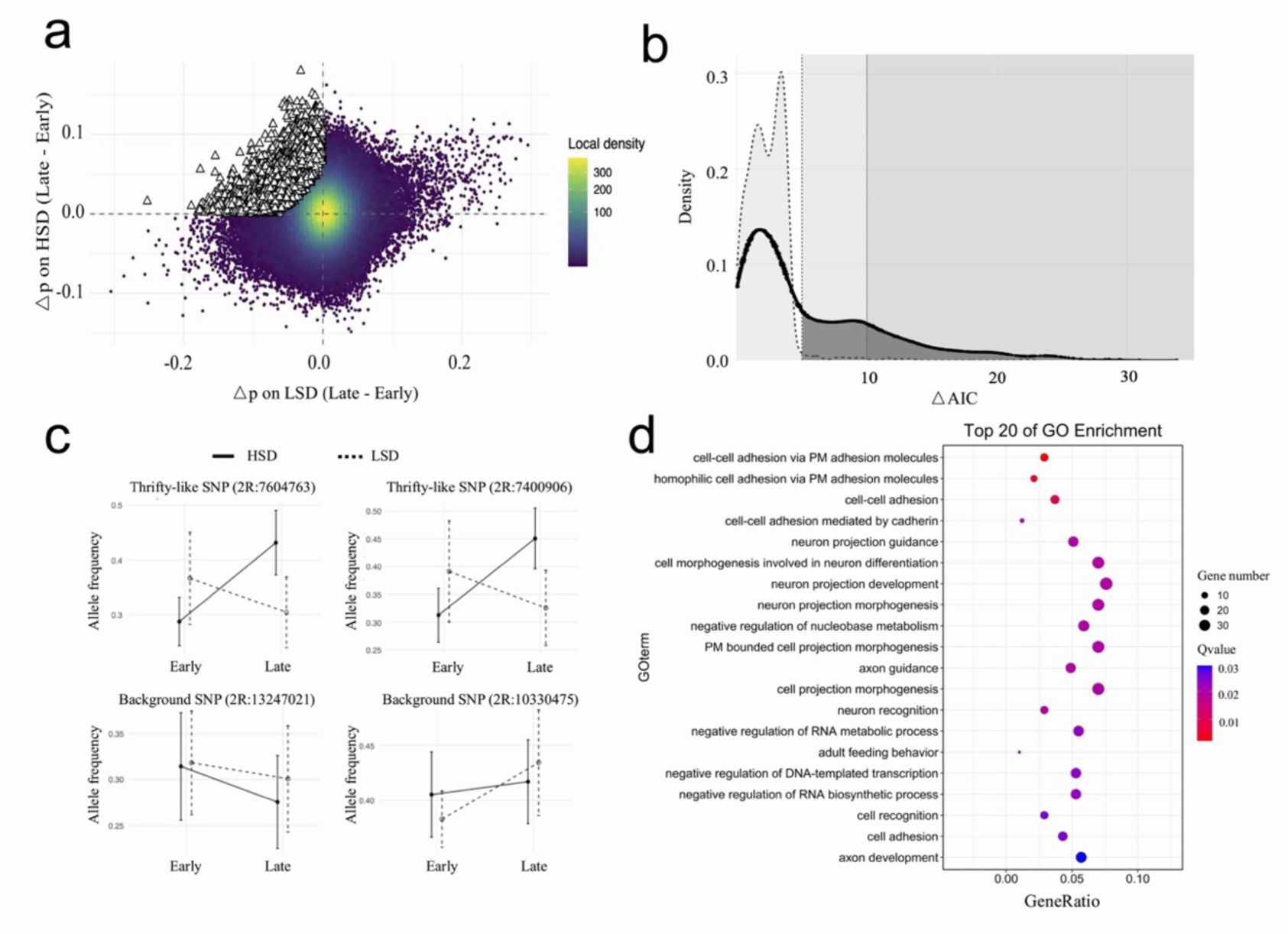
Identification and annotation of directional diet-opposed allele-frequency shifts associated with development time. **a**, Genome-wide comparison of allele-frequency shifts between early- and late-eclosing flies under LSD and HSD. For each SNP, allele-frequency shift Δp was calculated as p_Late_ − p_Early_ within each diet. The x-axis shows Δp under LSD, and the y-axis shows Δp under HSD. Point color indicates local SNP density, and open triangles indicate candidate thrifty-like SNPs, enriched in late-eclosing flies under HSD and depleted in late-eclosing flies under LSD. **b**, Model-based refinement of candidate thrifty-like SNPs. Density curves show ΔAkaike information criterion (ΔAIC) distributions for candidate thrifty-like SNPs and randomized background SNPs. The solid line indicates candidate thrifty-like SNPs, and the dashed line indicates the randomized background SNPs. The vertical dotted line marks the ΔAIC cutoff used to define robust thrifty-like SNPs. **c**, Representative allele-frequency patterns for thrifty-like and background SNPs. Allele frequencies are shown for early- and late-eclosing flies under HSD and LSD. Solid lines indicate HSD, and dashed lines indicate LSD, and error bars indicate s.e.. **d**, Gene Ontology (GO) biological process enrichment for genes associated with robust thrifty-like SNPs. The dot plot shows the top 20 enriched GO terms. The x-axis indicates gene ratio, dot size indicates the number of genes in each term, and dot color indicates the adjusted q value. PM, plasma membrane.

Representative SNPs illustrated the expected high-sugar delay-associated pattern. For example, SNPs such as 2R:7604763 and 2R:7400906 increased in frequency among late-eclosing flies under HSD but decreased among late-eclosing flies under LSD, suggesting that these alleles are specifically associated with delayed development in the high-sugar environment. By contrast, comparison SNPs such as 2R:13247021 and 3L:10330475 changed in the same direction under both diets (**Figure 5c**). Across the genome, many loci showed allele-frequency divergence between early- and late-eclosing individuals under both diets, but thrifty-like SNPs were distinguished by consistent opposite shifts across diets, with alleles enriched in late-eclosing flies under HSD tending to be depleted in late-eclosing flies under LSD.

Genes associated with these thrifty-like variants were significantly enriched for Gene Ontology (GO) biological process categories that converged on three related themes: cell adhesion, neural development, and morphogenesis (**Figure 5d, Supplementary Data 6**). The strongest terms included plasma-membrane and cadherin-mediated cell adhesion, neuron projection guidance and development, axon guidance, neuron recognition, and cell-projection morphogenesis. The broader enrichment set also included processes related to transcriptional regulation, feeding behavior, calcium and cyclic nucleotide signaling, synaptic target recognition, and appendage development. These findings suggest that thrifty-like variants may affect development time through adhesive, neural, and morphogenetic processes that connect dietary cues to growth and maturation decisions, rather than through core carbohydrate or lipid metabolic pathways alone.

### Candidate loci map to human metabolic genes

Finally, we asked whether the human homologs of our diet-responsive fly candidates have been linked to metabolic or anthropometric traits in humans (**Figure 6**). This comparison highlighted a subset of conserved genes with plausible roles in nutrient-responsive growth and metabolism. One of the strongest cross-species links was the G×E candidate *CG43373*, whose putative human ortholog *ADCY5* has been repeatedly associated with type 2 diabetes, fasting glucose, body mass index (BMI), height, and weight. Although *CG43373* remains poorly characterized in flies, its homology to a mammalian adenylyl cyclase suggests a potential connection to conserved cAMP-dependent regulatory pathways. Additional G×E candidates also mapped to human genes with metabolic relevance, including *Cerk*, *Pak*, *Eip75B*, and *Ac78C*, whose corresponding human orthologs have likewise been implicated in signalling or metabolic phenotypes.

**Figure 6.**
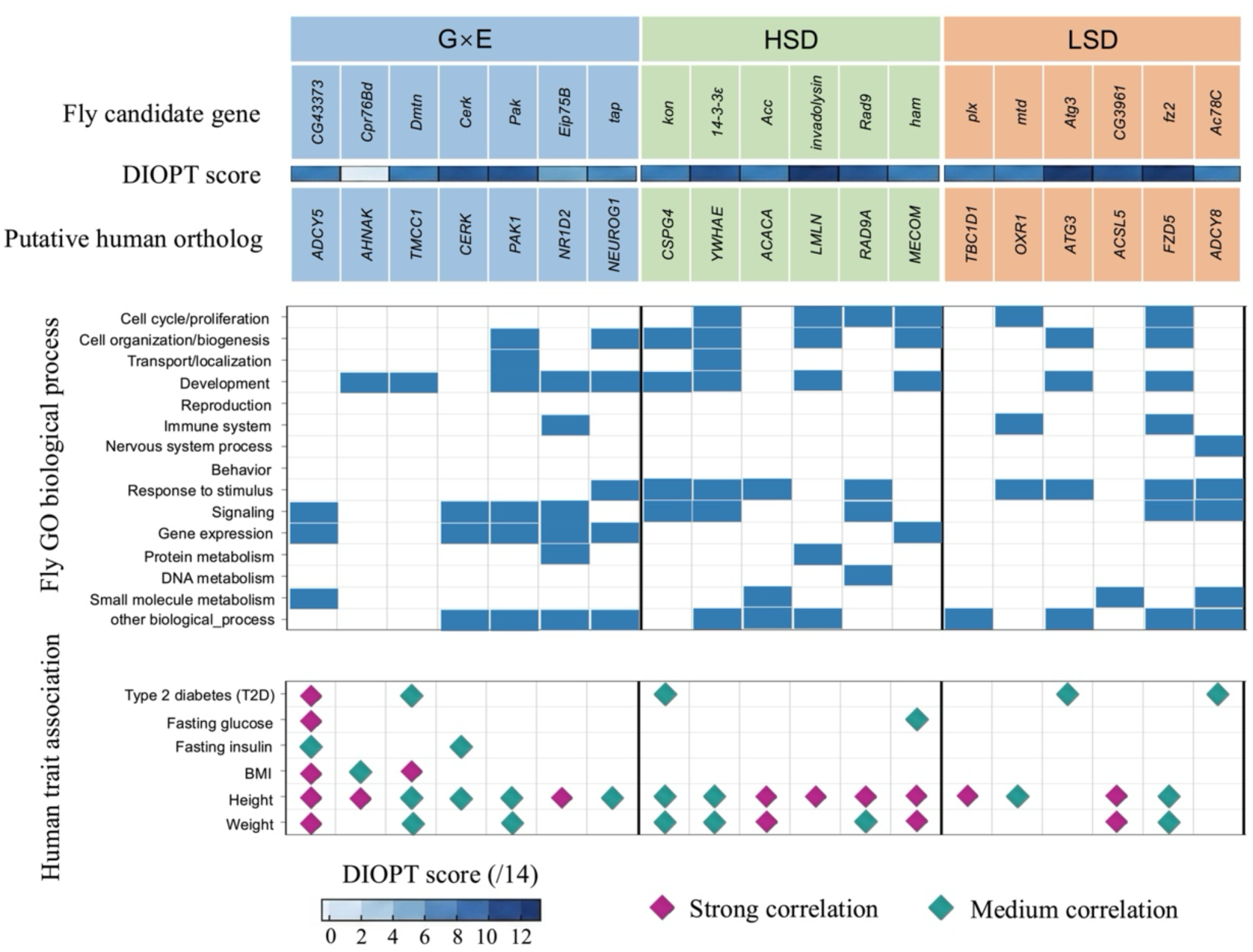
Cross-species annotation of fly candidate genes using human trait data. Candidate genes from the G×E, HSD and LSD analyses are grouped by discovery class. For each fly gene, the top rows show the putative human ortholog and DIOPT score, which summarizes support for the predicted fly–human ortholog relationship. The middle panel shows fly biological process annotations; blue boxes indicate annotated processes for each gene. The lower panel shows human metabolic or anthropometric trait associations reported for the corresponding human orthologs in the Type 2 Diabetes Knowledge Portal; diamonds indicate associated traits and colour denotes association strength. BMI, body mass index; T2D, type 2 diabetes.

We also identified notable human orthologs among diet-specific candidates. For example, *Acc* links a fly gene required for fatty acid biosynthesis to a central metabolic enzyme in mammals, whereas *Atg3* connects nutrient-dependent developmental phenotypes to the autophagy pathway, a key regulator of cellular energy balance. Similarly, *plx* is of particular interest because mammalian *TBC1D1* has been associated with body mass and glucose-related traits, consistent with a conserved role in nutrient-responsive physiology. Together, these comparisons suggest that, although not all fly candidates map to human trait-associated genes, a substantial subset falls within conserved pathways involved in signalling, lipid metabolism, autophagy, endocrine regulation, and growth control. These results highlight the utility of *Drosophila* genetics for nominating conserved nutrient-responsive loci for mechanistic follow-up.

## DISCUSSION

Dietary sugar does more than shift development time; it reshapes the genetic architecture underlying it. Using the DRPs, our newly developed outbred mapping system, we found that low- and high-sugar diets do not simply rescale genetic effects at a shared set of loci; instead, they alter which loci contribute to variation in development time, with additional loci emerging specifically through gene-by-diet interaction. This conclusion is consistent with prior *Drosophila* work showing that dietary composition can modify genetic and epistatic effects on development time^24^. Here, we extend this principle from predefined genotype panels to genome-wide mapping in outbred recombinant populations, allowing diet-specific and G×E loci to be resolved at the SNP and candidate-gene level. Consistent with this diet dependence, association signals were concentrated under LSD but more dispersed under HSD, suggesting that high-sugar stress recruits a broader set of loci contributing to development time. Functional validation further supports that several prioritized candidates act in a diet-dependent manner. Among the validated G×E candidates, *Eip75B* is especially notable because it encodes an ecdysone-inducible nuclear receptor and has previously been implicated in organismal sugar tolerance and high-sugar-induced de novo lipogenesis^23^. These functions link nutritional state to developmental progression, providing a plausible mechanistic bridge between the sugar environment and development time.

The DGRP provides a useful foundation for quantitative trait mapping because its sequenced inbred lines capture abundant natural variation and can be extended beyond fixed-panel association studies into outbred recombinant populations^16,17^. Several studies have since used DGRP-derived outbred recombinant populations to map the genetic basis of complex traits through large-scale phenotyping^25–27^. Based on this extensibility, our DRP framework combines known founder ancestry with additional recombination and experimentally controlled allele frequencies, thereby improving power to detect diet-dependent loci that may be rare or weakly represented in standard reference panels. First, each population starts from a defined subset of sequenced founders, which brings founder-derived alleles, including ones uncommon in the broader reference panel, to experimentally informative frequencies. Second, more than 30 generations of intercrossing accumulate recombination and break up founder haplotypes, improving mapping resolution. Third, paired populations sharing matched founder backgrounds provide an internal test of repeatability, so signals supported across populations are less likely to reflect population-specific noise. Coupled with extreme-phenotype sampling, pooled whole-genome sequencing, and Hidden Markov model (HMM)-based haplotype inference, this framework makes large-scale, cost-effective mapping of diet-dependent loci feasible in thousands of flies.

At a conceptual level, our results support a model in which nutritional environment reorganizes the genotype-to-phenotype map rather than merely modulating a fixed developmental program^28^. This is especially relevant for development time, which integrates growth, nutrient sensing, endocrine regulation, and stress adaptation. In *Drosophila*, conserved nutrient-sensing pathways, including insulin/IGF and TOR signaling, couple diet to growth and developmental progression^11,29^, and perturbation of insulin signaling is known to delay development^30^. Previous work showing that food type can alter development time and modify higher-order genetic interactions further supports the view that development time is highly sensitive to nutritional context^24^. Because development time reflects the cumulative coordination of these processes, it may provide a more informative systems-level readout of dietary challenge than individual metabolic endpoints alone. The broader DGRP phenotypic analysis supports this interpretation by showing that dietary sugar altered not only trait means but also the structure of phenotypic variation across genotypes, traits, and life stages. High sugar therefore does not simply alter trait values; it may also expose genetic and residual variation that is otherwise buffered under standard dietary conditions. This pattern is consistent with the concept of cryptic genetic variation, in which standing genetic variation that has little detectable phenotypic effect in one environment can become phenotypically relevant under novel or stressful conditions^31^. In this framework, development time emerges as an integrative trait that captures genotype-specific trade-offs among these processes.

The thrifty-like variants identified from directional diet-opposed allele-frequency shifts add a hypothesis-generating evolutionary perspective to these findings. These variants showed patterns consistent with conditional effects across nutritional environments, matching the defining pattern of alleles associated with relatively earlier development under lower-sugar conditions but delayed development under high-sugar stress. This pattern is conceptually related to thrifty-gene or nutritional mismatch models^5,7^, in which alleles that are neutral or beneficial under caloric scarcity may become disadvantageous under caloric or sugar excess. Because the classic thrifty genotype hypothesis remains debated^32,33^, our study does not resolve this evolutionary debate in humans. Instead, it provides a controlled experimental system to test whether alleles can have opposite phenotypic consequences under different sugar environments, and whether these diet-dependent developmental effects translate into context-dependent fitness consequences. Notably, the thrifty-like candidate set was enriched for cell-adhesion, neurodevelopmental and morphogenetic processes, rather than primarily for core carbohydrate or lipid metabolic pathways.

This functional profile suggests that diet-dependent developmental trade-offs may act through mechanisms that connect dietary cues to cell–cell communication, neural patterning, and growth and maturation decisions. The overlap between fly candidates and human genes associated with metabolic or anthropometric traits likewise suggests that at least part of this biology is conserved. Although these links do not imply one-to-one correspondence with human disease loci, they support the idea that nutrient-responsive variation repeatedly targets shared regulatory pathways relevant to metabolic regulation^1,34^.

Several limitations should be noted. First, although our results are consistent with diet-dependent or thrifty-like effects, direct tests of selection and fitness consequences across generations were beyond the scope of this study. Future experimental-evolution studies can test whether candidate alleles shift in frequency under different diets and whether these shifts are associated with viability, fecundity, or other fitness components. Second, functional validation supports roles for multiple candidates in diet-responsive development time, but these perturbation experiments prioritize candidate genes rather than resolving the causal natural variants or the specific tissues and pathways through which they act. Future allele-replacement, tissue-specific perturbation, and rescue experiments can define the causal variants and developmental contexts underlying these effects. Third, the sex-dependent validation results further suggest that some loci may act through sexually dimorphic mechanisms or higher-order sex × genotype × diet interactions. Future sex-aware mapping and functional assays can determine how sex modifies diet-dependent genetic effects on development time. More broadly, this work establishes the DRP framework as a tractable system for mapping context-dependent genetic effects and linking nutritional environment to developmental genetics.

## METHODS

### Fly husbandry and diets

To characterize diet-dependent phenotypic variation in founder backgrounds used for DRP construction, we measured a panel of metabolic and life-history traits in larvae and adults from 32 DGRP lines included among the DRP founders (lines #1–32 in **Supplementary Table 1**) reared on HSD or LSD. All experiments were conducted at 25 °C, 50% relative humidity, under a 12 h:12 h light:dark cycle. Flies were maintained on Bloomington standard cornmeal medium (http://flystocks.bio.indiana.edu/Fly_Work/media-recipes/bloomfood.htm). HSD and LSD were prepared from Bloomington semi-defined medium^35^, with sucrose concentration as the primary variable, following a previously established *Drosophila* high-sugar diet paradigm in which 1.0 M sucrose induces hyperglycemia, insulin resistance, lipid accumulation and developmental delay relative to a 0.15 M sucrose control diet^9^. Per liter, the diet contained agar (10 g), yeast (80 g), yeast extract (20 g), peptone (20 g), MgSO₄ (2 mL of a 1 M stock), CaCl₂ (3.4 mL of a 1 M stock), propionic acid (6 mL), and tegosept (10 mL of 10% w/v in 95% ethanol), supplemented with sucrose at 51.3 g (0.15 M; LSD) or 342 g (1.0 M; HSD). Unless otherwise noted, all assays were performed with five biological replicates. To minimize parental environment effects and larval crowding, flies were reared for two generations on standard medium under controlled density prior to experiments.

### Larval and adult metabolic phenotyping

We measured body weight, glucose, glycogen and triglyceride levels in pools of 10 larvae or 10 adult females from each line under both HSD and LSD treatments. One pool of 10 animals constituted one biological replicate. For the adult-stage experiments, flies were reared on standard medium from egg-to-adult eclosion under controlled larval density for each line. Two-day-old sexually mature adult flies were then transferred to HSD or LSD for 5 days at a density of 50 males and 50 females per bottle, with transfers to fresh food every 2–3 days. Adult females were collected for body weight and metabolite assays. For the larval stage experiments, animals were raised on HSD and LSD from embryos to third-instar stage (L3). Wandering L3 larva were staged by adding blue food dye to the food: larvae with blue guts were classified as early wandering L3, while larvae with clear guts were classified as late wandering L3 and not used. Early wandering L3 larvae were collected for body weight and metabolite assays.

Each group of 10 animals was weighed and then homogenized in 100 μL PBS buffer (Roche) containing 0.05% Tween at 4°C using a Geno Grinder (Spex Sample Prep) at 1,700 strokes/min for 1 min with three 2.3 mm grinding balls. Whole-body metabolite assays were performed using standard approaches for *Drosophila* metabolic measurements described previously^36^. The homogenate was heated at 70°C for 10 min to inactivate endogenous enzymes, and the heat-inactivated supernatant was used for glucose, glycogen, and triglyceride assays. Free glucose levels were measured using Hexokinase Reagents (Thermo Scientific, TR15421). Glycogen was digested with amyloglucosidase from Aspergillus niger (Sigma-Aldrich, A7420) and measured by subtracting free glucose from total glucose. Triglyceride level was quantified using Infinity Triglyceride Reagent (Thermo Scientific, TR22421). Glucose and glycogen standards were processed in parallel and used to quantify the sugar levels in the samples. All assays were conducted following the manufacturer’s instructions, and the results were normalized by the weight of each sample.

To compare whole-body and circulating glucose measurements, hemolymph glucose levels in adults were measured in eight lines by puncturing the thorax with a tungsten needle and then collecting the hemolymph by centrifuging the punctured flies through a silica membrane^36^. We compared glucose measurements from whole-body homogenate and hemolymph preparations. Fold changes in glucose levels (HSD/LSD) showed a consistent trend between the two sample types (**Supplementary Figure 1**). The homogenate method required only about one fifth of the flies required for the hemolymph method and showed greater consistency among replicates, as indicated by a smaller standard error of the mean. We therefore used whole-body glucose measurements for the remaining lines.

### Life-history trait phenotyping

We measured three life-history traits under HSD and LSD: egg-to-pupa development time, larval survivorship from first instar larvae to pupae, and adult longevity. Embryos were collected for each line by allowing fertilized females, 2-5 days post eclosion, to lay eggs for 3 h; L1 larvae were collected 24 h after egg deposition and placed on high or low sugar diets, 70 individuals per vial. White pupae formed on the vial walls were counted every 12 h to generate cumulative pupation curves and the total number of pupae was also used to estimate larval survivorship. Development time curves were generated by fitting a sigmoid function to the cumulative pupation data. The code is available on Github (https://github.com/xuanzhuang/DRP/tree/main/sigmod_fitting). Development time was estimated from these curves as the number of hours from egg deposition to when 50% of the total pupae had formed. Virgin females were reared on standard diet for two days following eclosure and then transferred to HSD and LSD, five age-matched individuals per vial and five replicates per diet treatment. Flies were then transferred to fresh medium every two days without using CO_2_ and the numbers of live flies in each line were recorded until all were deceased. The median lifespan was recorded for each line.

### Trait variance heritability and correlations

Larval and adult traits were measured for 32 DGRP lines under both HSD and LSD conditions. Data were collected from five biological replicates per line per treatment. Statistical analyses were carried out using R version 4.1.1. The distribution of each trait was examined separately for each life stage and diet. The relative change in each trait for each line between HSD and LSD was calculated from line means for each diet as (HSD − LSD)/LSD, and standard error (SE) of the relative change was calculated using the mean and variance of each diet as follows:

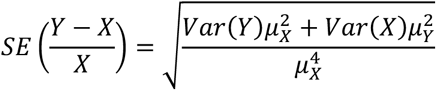

where *Y* represents HSD and *X* represents LSD. Data were then analyzed using two-way ANOVA with replication. DGRP lines (#1–32) and treatments (HSD or LSD) were used as independent variables, with measured traits as dependent variables.

Variance and heritability were analyzed using the R package variability. Within-line variance (V_E_) was calculated from the five biological replicates of each line for each treatment, while between-line variance was calculated and used as a proxy for Genetic Variance (V_G_). Total Phenotypic Variance (V_P_) = V_G_ + V_E_. Broad sense heritability was calculated as the proportion of V_G_ in V_P_. The genotypic coefficient of variance (CV_G_) and residual coefficient of variance (CV_R_) were calculated by standardizing V_G_ and V_E_, respetively, by the trait mean X^37^.

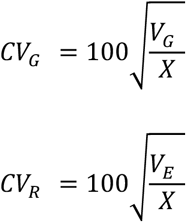

Pairwise phenotypic trait correlations were calculated in R using the Spearman method. Three sets of data were analyzed: relative phenotypic change between two dietary treatments (HSD − LSD)/LSD, phenotype on HSD, and phenotype on LSD. In addition, we also analyzed the correlation between HSD and LSD treated phenotypes of each trait. Line mean values were used to determine the correlation coefficients and P-values were calculated for each correlation.

### DRP construction and mapping design

We developed a multi-parental advanced intercross mapping resource, termed the DRPs, to map natural genetic variation affecting development time under different dietary sugar conditions (**Figure 1**). The DRP comprises 16 independently maintained AIPs founded from a total of 64 inbred *Drosophila* Genetic Reference Panel (DGRP) lines (**Supplementary Table 1**). Each AIP was initiated with eight founder DGRP lines, and the 64 founders were allocated such that each line contributed to two AIPs, enabling cross-population validation of mapped signals. To reduce potential bias introduced by mating preference and to minimize viability differences among founder lines during establishment, AIPs were generated through two generations of defined reciprocal round-robin crosses among the eight founders, followed by pooling of offspring to seed each synthetic population. After population founding, flies were maintained under random mating for ≥30 generations (typically 30–40 generations) to accumulate extensive recombination. Each AIP was maintained at a census size of approximately 5,000–7,000 individuals, thereby minimizing genetic drift while preserving a highly heterogeneous and recombined mapping population.

Discrete generations were propagated by collecting eggs from fertilized females over approximately 24 h and distributing them into 24 bottles per AIP (approximately 200–250 eggs/bottle). Bottles were then placed into a cage with bottle tops removed so that newly eclosed virgin adults could freely encounter one another, ensuring random mating and initiation of the next generation. For dietary treatments, we used HSD and LSD formulations based on the same basic recipe. To improve comparability of egg-to-adult survival between diets and maximize the number of individuals contributing to development time measurements, the volume of water added to each recipe was fixed at 900 mL for both HSD and LSD. For each AIP, embryos were collected and reared on each diets (10 bottles per diet per population). Development time in the DRP assay was indexed by eclosion time, and we implemented an extreme-phenotype mapping strategy. Within each AIP and diet, we selected individuals from the earliest 5% and latest 5% of eclosing flies. For each diet and population, we collected 96 females from each phenotypic extreme. In total, 6,144 flies were sampled for genetic mapping. Selected individuals were individually barcoded and pooled for whole-genome sequencing, enabling parallel genome-wide association analyses for HSD, LSD, and a combined G×E model to identify loci with diet-dependent effects on development time.

### DNA extraction and individually barcoded genome sequencing

Genomic DNA was extracted from individual flies in a 96-well format using a bead-based kit (Agencourt DNAdvance, Beckman Coulter), with all reagent volumes reduced to one quarter of the manufacturer’s recommended protocol. DNA concentrations were quantified using the Quant-iT PicoGreen dsDNA assay Kit (Invitrogen). All 6,144 samples were individually indexed with the Nextera 96 Sample Index Kit (Illumina, FC-121-1012), and sequencing libraries were prepared using the Nextera DNA Flex Library Prep Kit (Illumina, FC-121-1031) with a miniaturized protocol that reduced reaction volumes (following an approach similar to Baym et al.^38^). Indexed libraries were pooled in sets of 96 for sequencing. To ensure approximately equal representation of each individual within a pool, libraries were quantified and normalized prior to pooling. Sequencing was performed at the University of Chicago Genomics Facility on an Illumina HiSeq 4000 platform using 150-bp paired-end reads (https://voices.uchicago.edu/genomicsfacility/).

### Sequence assembly and SNP calling

Short-read data were aligned to the *Drosophila* reference genome (v6; http://flybase.org) using the BWA-MEM alignment algorithm implemented in BWA v0.7.15^39^ with default settings. Read coverage across the reference genome was summarized using SAMtools mpileup^40^, and the output was used for variant calling using BCFtools v1.4.1^41^. Sites with total coverage ≥ 500 or read-mapping quality scores ≤ 20 were excluded from downstream analyses. Only the major chromosomal arms 2L, 2R, 3L, 3R, and X were retained for analysis. Chromosome 4 was excluded because it comprises only a small proportion of the *Drosophila* genome, contains interspersed heterochromatic and euchromatic DNA, and does not undergo crossing over^42^.

### HMM founder haplotype imputation

We employed a HMM to impute the founder haplotypes for each sequenced fly from low-coverage sequence data and the available founder genomes. Each genotype of a DRP individual is produced by a pair of recombinant chromosomes derived from founder haplotypes, referred to as the founder diplotype. The purpose of the HMM is to infer the mosaic founder diplotype underlying each genome. The HMM we developed is a modified version of the HMM framework and Perl code of King et al^43^. To make the computation more efficient for our dataset, it is written in C; the complete code is available on GitHub (https://github.com/xuanzhuang/DRP/tree/main/HMM).

The HMM computes the probability of each possible founder ancestry state at each SNP site. Each AIP of the DRPs was seeded by eight parental founder lines, so there are 36 possible founder states: eight homozygous states, in which a locus is derived from the same founder on both chromosomes, and the 28 heterozygous states, in which two different founder haplotypes are present:

[utbl1]

The HMM consists of a Markov chain of hidden states and a set of observed variables, and each observation depends only on the underlying hidden state. In this HMM, the observations are the observed allele counts at each sequenced genomic position, and the hidden states are founder ancestry states. The model requires three sets of parameters: (1) initial probabilities: the prior probability for each possible founder ancestry state; (2) emission probabilities: the probability of observing each allele given the founder state at each locus; and (3) transition probabilities: the probability of transitioning from the founder state of one locus to that of the adjacent locus.

If we assume random mating among the eight founder lines in each AIP of the DRPs, and no selection occurs during the DRP construction, the initial probabilities are set according to the expected frequencies of founder states in the above matrix, including the 28 heterozygous states in the lower triangle, which are not shown. each homozygous state = 1/64 each heterozygous state = 1/32 Transition probabilities were described in two categories based on homozygous or heterozygous states and each category was further divided into four or five subcategories based on types of allele change (**Supplementary Table 2**). The transition probability is governed by the probability *r* of a recombination event occurring in the interval of two adjacent loci. These probabilities depend on the chromosome recombination rate occurring between two adjacent loci and the number of generations following DRP creation. In our model, the transition probabilities were calculated from an LD-based estimation of recombination rates^44^ as reported by the program LDhelmet. Recombination rates at positions between the observed markers were estimated by linear interpolation.

The emission probabilities were calculated from the observed allele counts, including alternative-allele and total-allele counts, using binomial distributions described in King et al.^43^. Parameterized by these three sets of probabilities, the HMM employs a standard forward and backward algorithm to calculate the probabilities of each possible founder state at each locus for each sequenced fly.

The reconstruction of an individual haplotype can be obtained by assigning each locus to the founder diplotype state that has the highest probability; only loci with this probability > 0.95 were retained. The diplotypes are phased to obtain a pair of haplotypes by minimizing the number of recombination events.

### Genome-wide association mapping

We performed quality control of genotype data by excluding SNPs with all missing genotypes and minor allele frequency (MAF) < 0.05 using PLINK v1.90^45^. To reduce redundancy among markers in high linkage disequilibrium (LD), SNPs were subsequently pruned using the PLINK option --indep-pairwise 100 1 0.8, which evaluates pairwise LD within a sliding window of 100 SNPs, advances the window by one SNP at a time, and removes one SNP from each pair with r² > 0.8. This threshold was used to remove highly correlated markers while retaining informative variation. To account for population structure and relatedness, we first estimated the genetic relationship matrix (GRM) using GCTA v1.92.2^46^, and incorporated the GRM as a random effect in downstream association analyses.

GWAS were conducted using GEMMA. We performed (i) diet-specific GWAS for HSD and LSD and (ii) genome-wide G×E analysis. For the diet-specific analyses, each individual AIP (DRP population) was analyzed separately using a linear mixed model including the GRM.

To test for GxE interactions between SNPs and diet (HSD vs. LSD), we fitted the model:

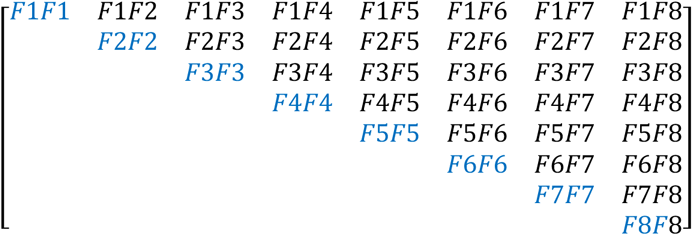

where *y* is the development time phenotype, coded as 0 for the early-eclosing tail and 1 for the late-eclosing tail, *G* is the genotype for a SNP, *E* is the diet environment (HSD or LSD), and *e* is the residual error. For the G×E analysis, we filtered out SNPs with missing rate > 0.9.

We then performed meta-analysis across individual AIPs by combining GWAS results using METAL^47^. To further reduce false positives, especially for low-frequency variants, we leveraged the founder-overlap structure of the DRP. As shown in **Figure 3a**, each founder contributes to two AIPs, creating pairs of populations that share a founder while otherwise representing independent mapping populations. Under this design, associations driven by founder-derived variants are expected to recur in AIPs that share that founder, whereas associations driven by AIP-specific stochastic effects (e.g., drift or segregation noise) or technical artifacts are less likely to do so. Therefore, we applied a design-informed replication filter and retained only loci supported in at least two AIPs (N ≥ 2), including one pair of AIPs that share a founder. This criterion prioritizes signals that are reproducible across founder-overlapping AIPs and reduces the contribution of AIP-specific artifacts.

### Polygenic score analysis

PGS analysis was performed as a supplementary analysis to assess whether the cumulative effects of GWAS-associated variants captured development time variation between early- and late-eclosing flies within each DRP population. The analysis was conducted separately for HSD and LSD, and separately for each of the 16 AIPs. For each diet and cage, diet-specific GWAS results generated by GEMMA were used as the source of SNP effect sizes. Genotype files after LD pruning were converted to additive allele-dosage format using PLINK v1.90.

For each GWAS result file, SNP identifiers were harmonized with the additive genotype matrix using the marker ID and effect allele. SNPs were then selected according to a series of GWAS *P* value thresholds: 1.0, 0.9, 0.8, 0.7, 0.6, 0.5, 0.2, 0.1, 0.05, 0.01, 0.005, 0.001, 0.0005, and 0.0001. At each threshold, only SNPs that were present in both the GWAS summary statistics and the genotype dosage matrix were retained. This threshold series was used to test whether the association between the PGS and phenotype increased as broader sets of GWAS-ranked SNPs were included.

For each individual, the PGS was calculated as the weighted sum of additive effect-allele dosages, using the GEMMA *β* estimate as the SNP weight. Thus, for individual i, the score was calculated as:

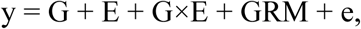

where 𝐺_ij_ is the additive dosage of the effect allele at SNP 𝑗, 𝛽_j_ is the corresponding GEMMA-estimated SNP effect size, and 𝑚is the number of SNPs included at a given *P* value threshold.

For each diet, AIP, and *P* value threshold, individuals with complete genotype and phenotype data were retained, and early-versus-late eclosion status was regressed on the polygenic score. The coefficient of determination (𝑅^2^) from this model was used to summarize the in-sample explanatory performance of the score within each AIP × diet dataset.

### Thrifty-like SNP identification

Genotype data from individually barcoded early- and late-eclosing flies were used to quantify allele-frequency changes within each AIP under HSD and LSD. For each SNP, allele frequency within each AIP × diet × developmental group (early or late) was calculated from the summed allele dosages of individual flies. A single counted allele was then used consistently across all AIPs and diets; when allele coding differed across records, frequencies were flipped so that allele-frequency changes always referred to the same allele.

Within each AIP and diet, we computed the developmental allele-frequency shift as Δp = pLate − pEarly, where pLate and pEarly denote the allele frequencies in late- and early-eclosing flies, respectively. SNPs were excluded from an AIP × diet comparison when both early- and late-eclosing groups were effectively monomorphic for the same allele, because such comparisons provide little information about allele-frequency shifts.

Sampling uncertainty for Δp was approximated from the finite number of sampled alleles in the early- and late-eclosing groups, corresponding to up to 192 alleles from 96 diploid flies per group when all individuals were successfully genotyped. For each SNP, informative AIP-level Δp estimates were then combined separately within each diet using inverse-variance weighting, generating two diet-specific summary shifts, ΔpHSD and ΔpLSD, with corresponding standard errors. Diet dependence was assessed by testing the directional diet contrast, ΔΔp = Δp_HSD_ − Δp_LSD_, against zero using a two-sided test.

We next screened for SNPs that showed a specific directional diet-opposed pattern in developmental allele-frequency shifts. In this study, we operationally defined these SNPs as “thrifty-like variants”, representing alleles enriched in late-eclosing flies under HSD but depleted in late-eclosing flies under LSD. A SNP was retained as a candidate thrifty-like variant if it met all of the following criteria (**Supplementary Figure 8**): informative comparisons in at least four AIPs under each diet; opposite directions of allele-frequency change between diets, with Δp_HSD_ > 0 and Δp_LSD_ < 0, corresponding to alleles enriched in late-eclosing flies under HSD but depleted in late-eclosing flies under LSD; a minimum directional diet contrast, ΔΔp = Δp_HSD_ − Δp_LSD_ ≥ 0.04; statistical significance of this contrast after multiple-testing correction, FDR < 0.10; and directional consistency in at least 70% of informative AIPs under each diet.

To further refine the candidate set, we used a binomial generalized linear model framework to test whether each SNP was best explained by a diet-dependent difference between early- and late-eclosing flies. The number of reference alleles was modeled out of the total number of sampled alleles across individually genotyped flies using three alternative models: an intercept-only null model, A0: ∼ 1; an additive main-effects model, A1: ∼ grp + diet; and an interaction model, A2: ∼ grp * diet, equivalent to grp + diet + grp:diet. Here, grp represents developmental group, early versus late eclosion, and diet represents HSD versus LSD. The interaction model therefore tests whether the allele-frequency difference between early- and late-eclosing flies depends on diet.

For each SNP, the best-supported model was defined as the model with the lowest Akaike information criterion (AIC) among A0, A1, and A2. ΔAIC was calculated as AIC_second-best_ − AIC_best_, so that larger ΔAIC values indicate stronger support for the best-fitting model. Candidate thrifty-like variants were then evaluated against a null distribution generated from randomly sampled background SNPs that did not pass the candidate filters described above. When more than 1,000 background SNPs were available, 1,000 SNPs were randomly selected. For each sampled background SNP, developmental-group labels were randomly permuted within each AIP × diet stratum, while preserving the original AIP identity, diet assignment, allele-count structure, and group sample size. The same model-fitting procedure was then applied to the randomized data to calculate ΔAIC values for the null distribution (**Figure 5b**). We classified SNPs for which the interaction model was best supported and ΔAIC > 5 as robust thrifty-like variants, indicating reproducible diet-dependent allele-frequency differences between early- and late-eclosing flies with the expected HSD late-enriched and LSD late-depleted pattern.

### GO enrichment analysis

To identify whether genes associated with candidate SNPs were enriched for particular biological processes, we performed GO enrichment analysis in *Drosophila*. Analyses were performed separately for each candidate SNP set. Candidate SNPs were assigned to nearby genes based on overlap with annotated gene bodies within a ±5 kb window around each SNP, using FlyBase annotation release r6.63. The resulting nonredundant gene lists were analyzed with the R package clusterProfiler using the *Drosophila* annotation database org.Dm.eg.db, with all annotated genes in the FlyBase r6.63 GTF used as the background set. *P* values were adjusted for multiple testing using the Benjamini–Hochberg method.

### Candidate gene validation

We selected 18 candidate genes for functional validation in *Drosophila* using transgenic UAS-RNAi knockdown (KD), UAS overexpression (OE), and independent loss-of-function (LOF) lines from the Bloomington *Drosophila* Stock Center (http://flystocks.bio.indiana.edu) (**Supplementary Data 7**). We used a standardized phenotyping workflow (**Figure 4a**): for UAS-based assays, candidate responder lines were crossed to a ubiquitin-GAL4 driver, and egg-to-adult development time was compared between F1 progeny carrying the intended perturbation and sibling control genotypes from the same cross. In parallel, LOF candidates were assayed using independently generated LOF lines together with genetic background–matched LOF control lines.

For the development time assay, embryos were obtained from 300–500 pairs of sexually mature flies for each cross within a 2 h collection window. Embryo density was tightly controlled and matched across diets and genotypes to minimize confounding effects of larval crowding and nutrient competition. Approximately 200 embryos were distributed into each of three bottles containing either LSD or HSD. After the first adult emerged, newly eclosed adults were collected every 2 h until eclosion ceased. Flies were sorted and counted according to sex and visible marker phenotypes, including eye color and wing morphology, which were used to assign each individual to the perturbed or matched-control class. Egg-to-adult development time for each individual was defined as the duration between the midpoint of the egg collection window and the midpoint of the adult sampling interval in which the individual eclosed.

Statistical analyses were performed separately for males and females. Within each sex and diet, we compared development time between perturbed and control individuals for each candidate gene using a two-sided Wilcoxon rank-sum test. To evaluate diet-dependent effects (G × E), we fitted a linear model for each candidate gene:

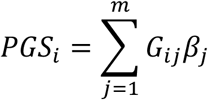

where perturbation status indicates individuals carrying the intended perturbation versus matched-control individuals. The perturbation status × diet interaction was assessed by ANOVA.

### Cross-species functional annotation

To functionally annotate candidate genes, *Drosophila* genes identified from the diet-responsive and G×E analyses were annotated in FlyBase to summarize known or predicted biological functions and associated pathways. For cross-species comparison, each *Drosophila* candidate gene was mapped to its human orthologs, and the human gene with the highest DIOPT score was selected as the representative putative human ortholog for downstream annotation^48^.

To assess whether these human orthologs were associated with metabolic or anthropometric traits, we queried the Type 2 Diabetes Knowledge Portal (T2D Knowledge Portal; http://www.type2diabetesgenetics.org/). Reported trait associations were recorded for each selected human ortholog, with emphasis on type 2 diabetes, glucose-related traits, BMI, body weight, height, lipid-related traits, and other metabolism-related phenotypes. The resulting annotations were used to summarize potential functional conservation between *Drosophila* candidate genes and human metabolic trait-associated loci.

## Acknowledgements

We thank Matthew Stephens for advice on HMM-based founder-haplotype imputation, Graeme Bell for valuable conceptual suggestions, Gabriella Ceresa and Hengxing Zou for experimental assistance, and Natalia Tamarina for helpful feedback on the manuscript. We acknowledge the Arkansas High Performance Computing Center which is funded through multiple National Science Foundation grants and the Arkansas Economic Development Commission. This work was supported by the National Institute of General Medical Sciences of the National Institutes of Health under award numbers R35GM160135 and R15GM152956 to X.Z., R01GM114289 to M.K., and P20GM139768 through the Arkansas Integrative Metabolic Research Center COBRE program supporting X.Z.’s Research Project Leader subproject; by the Arkansas Biosciences Institute to X.Z.; and in part by the National Institute of Diabetes and Digestive and Kidney Diseases of the National Institutes of Health under award number P30DK020595.

## Author contributions

X.Z. and M.K. conceived the study. X.Z. supervised and directed the project. X.Z. and Y.B. designed the experiments and analyses. X.Z., M.A., and Y.B. generated and analyzed the DGRP phenotypic data. M.L., S.P., and X.Z. developed and maintained the DRP populations. X.Z. performed the DRP phenotyping and sample collection, prepared sequencing libraries, and developed the HMM-based founder-haplotype inference pipeline. F.M. and Y.I.L. performed the initial raw sequencing-data processing and preliminary GWAS analyses. Y.B. performed the final GWAS and G×E analyses, developed the DRP built-in replicate-aware post-GWAS screening strategy, and performed the allele-frequency, GO-enrichment, cross-species, and functional validation analyses. Y.B., S.S., Y.C., S.A., A.T., and M.R. performed the functional validation experiments. Y.B. generated figures and tables. Y.B. and X.Z. interpreted the results and wrote the manuscript. All authors reviewed and approved the final manuscript.

## Data availability

The authors declare that the main data supporting the findings of this study are available within the article and its Supplementary Information files. Extra data are available from the corresponding author upon request.

