## Supplementary Tables and Figures for "Outbred *Drosophila* populations reveal diet-dependent genetic effects on development time": Description of Additional Supplementary Files.docx

File Name: Supplementary Data 1.

Description: Phenotype summary and line-level data for 32 DGRP lines under LSD and HSD conditions, including diet responses, missing-line information, *P* values, and correlation matrices.

File Name: Supplementary Data 2.

Description: SNPs exceeding different nominal significance thresholds in meta-GWAS analyses under HSD, LSD, and G×E models.

File Name: Supplementary Data 3.

Description: Pilot validation assay counts from responder stock × driver #32551 crosses under LSD and HSD.

File Name: Supplementary Data 4.

Description: Summarized development time records from candidate-gene validation assays.

File Name: Supplementary Data 5.

Description: List of 1082 identified candidate thrifty-like SNPs with ΔAIC values.

File Name: Supplementary Data 6.

Description: Top 30 enriched GO biological process terms for Thrifty-like SNPs and development time-associated SNPs identified under HSD, LSD, and G×E models.

File Name: Supplementary Data 7.

Description: *Drosophila* stocks used for functional validation of candidate GWAS genes.
