## Supplementary Tables and Figures for "Outbred *Drosophila* populations reveal diet-dependent genetic effects on development time": Supplementary Information.docx

**Zhuang et al.**

**SUPPLEMENTARY INFORMATION**

Supplementary Figures

#### Supplementary Figure 1. Comparison of glucose measurement from whole-body homogenate and hemolymph.

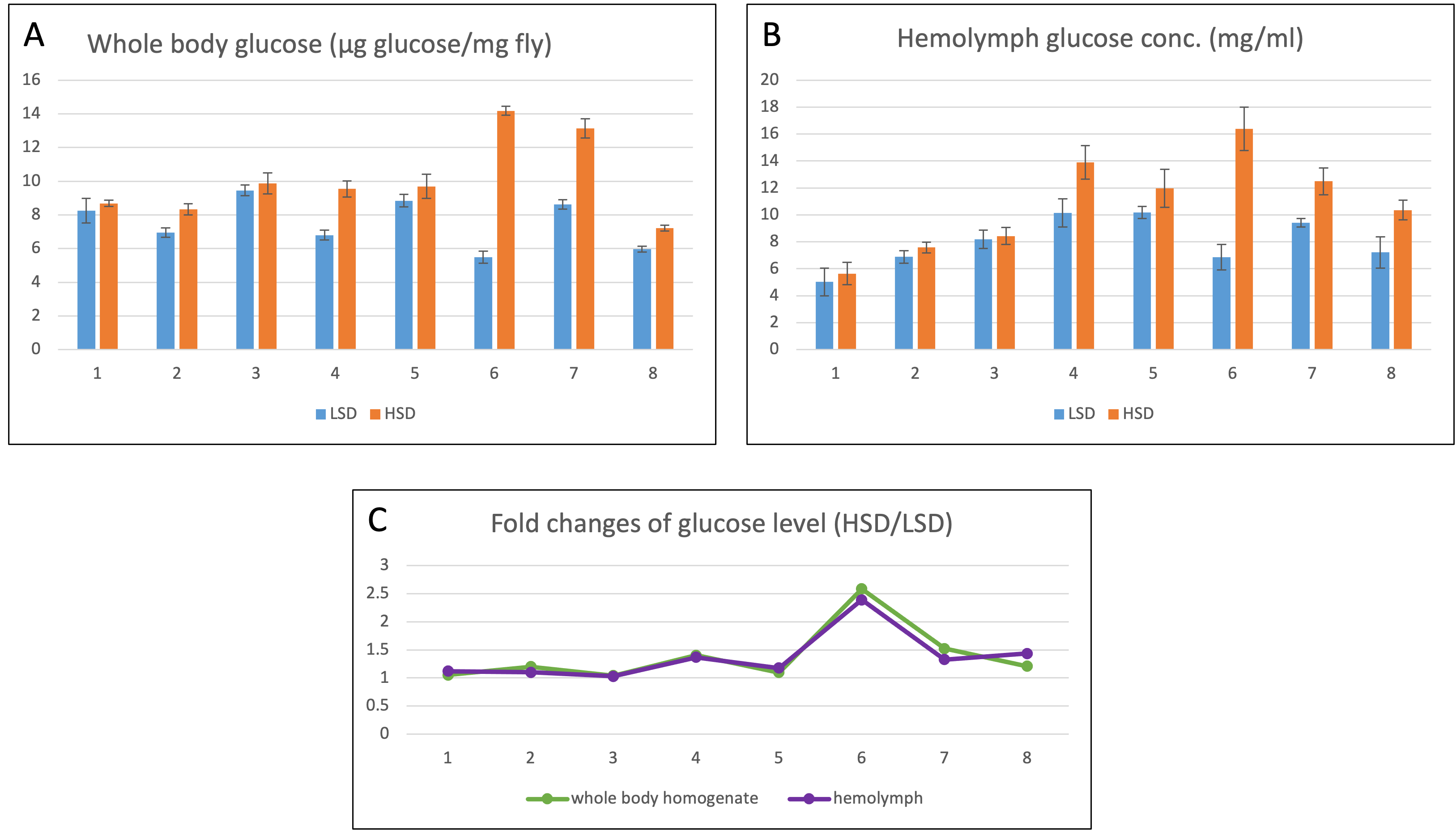

(A) Glucose level of whole-body homogenate, normalized by body weight. (B) Hemolymph glucose concentration. (C) HSD/LSD fold changes of glucose level, comparing the trend of the eight lines in A and B. Y axis indicates glucose level (A and B) or fold change of HSD/LSD (C), and X axis indicates the first eight lines (#1-8) used in the study (see Table S1 for DGRP line information). Error bar in A and B represent the standard error of the mean of five replicates.

#### Supplementary Figure 2. Q-Q plots of (A) HSD, (B) LSD, and (C) G × E GWAS.

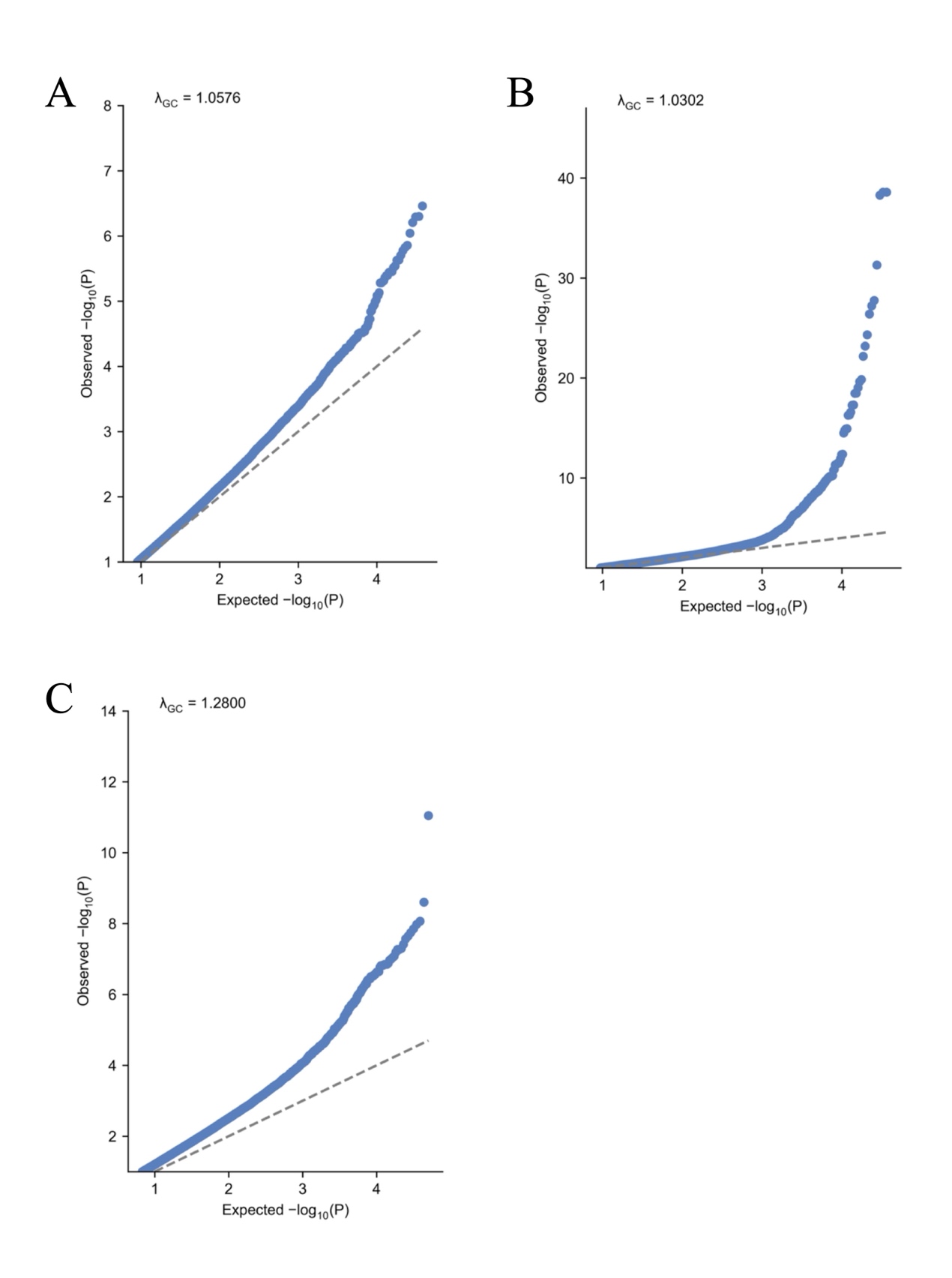

#### Supplementary Figure 3. Venn diagram of SNPs exceeding the nominal significance threshold in the HSD, LSD, and G × E GWAS.

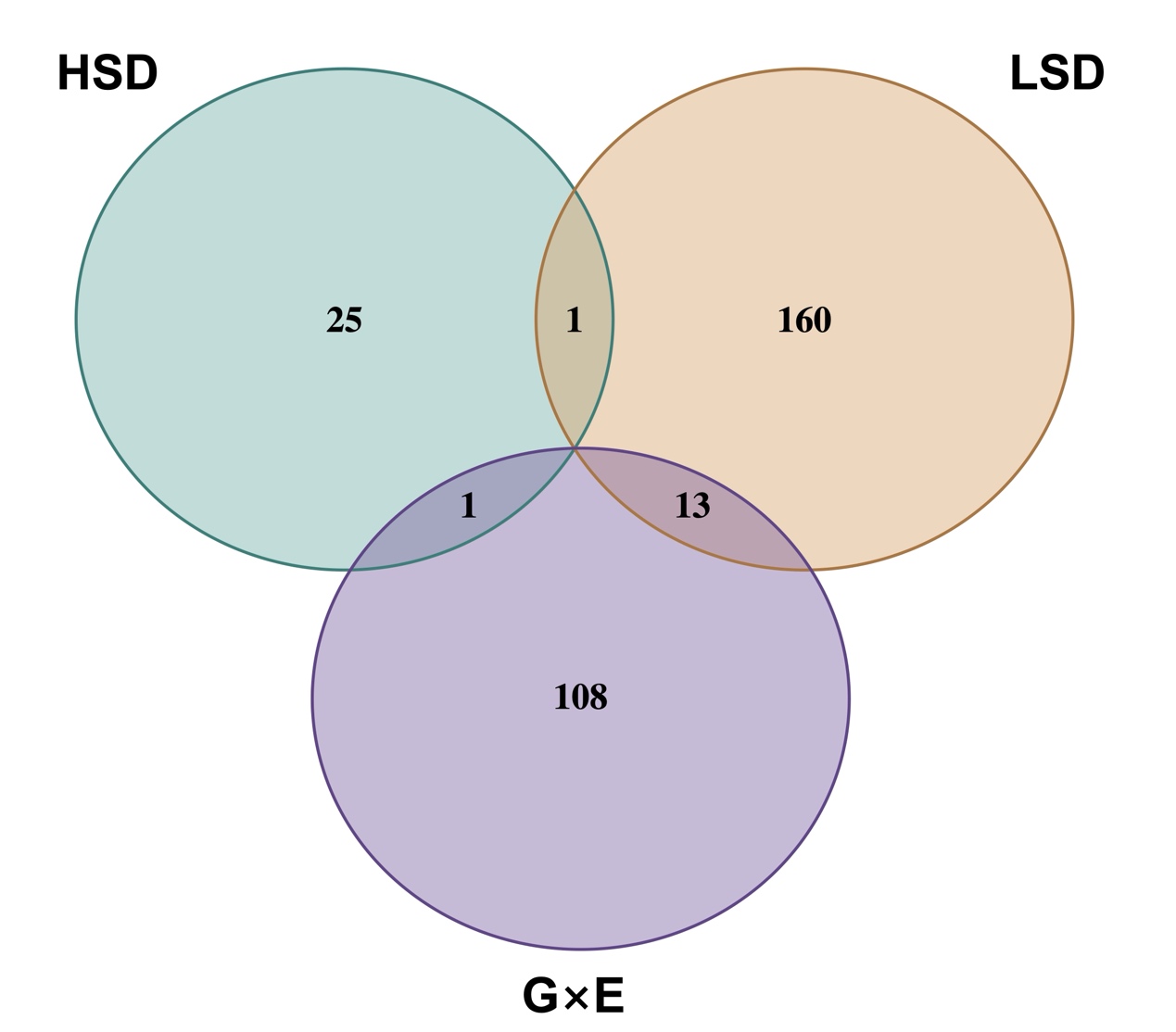

SNPs surpassing the nominal significance threshold (P < 1 × 10⁻⁵) in each GWAS were compared, and the numbers of overlapping SNPs between analyses are indicated in the shared regions of the corresponding circles.

#### Supplementary Figure 4. Manhattan plots of GWAS results for each AIP (cage) under (A) HSD and (B) LSD.

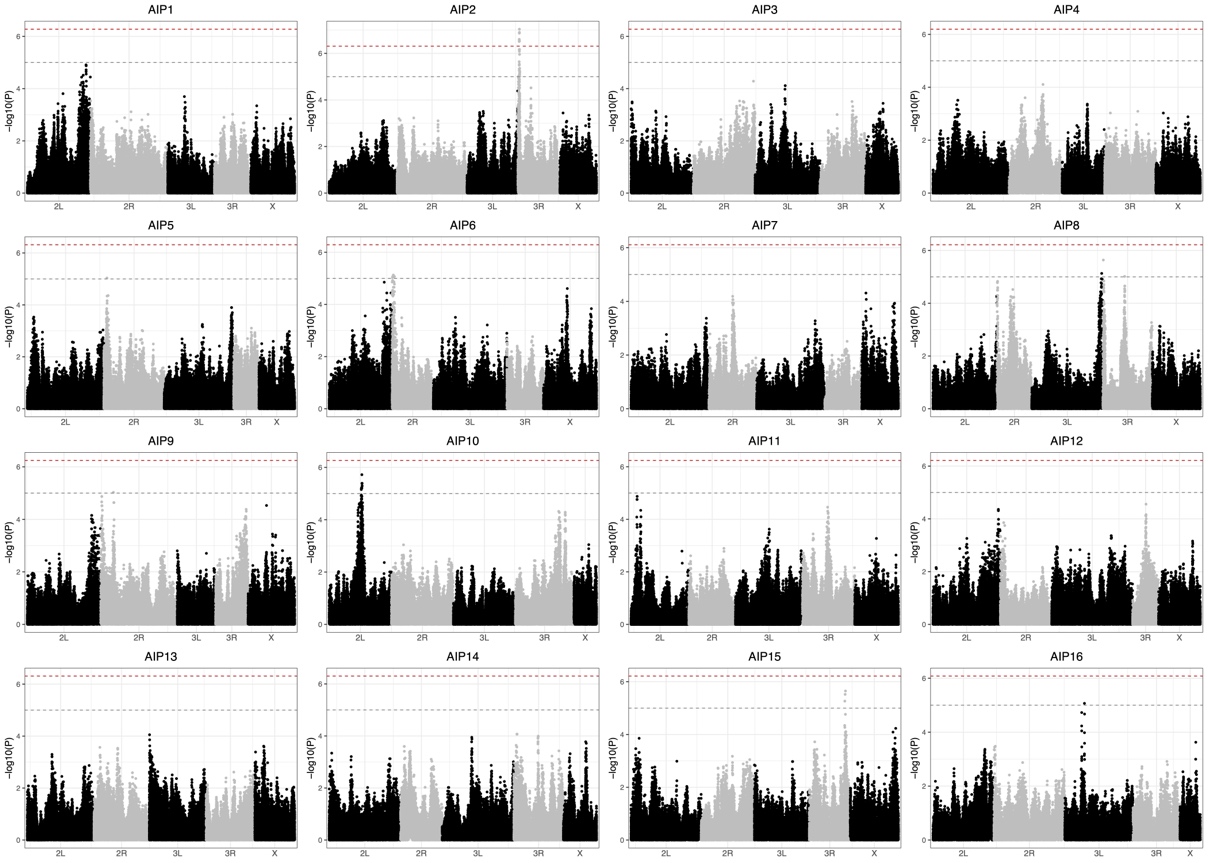

（A）HSD

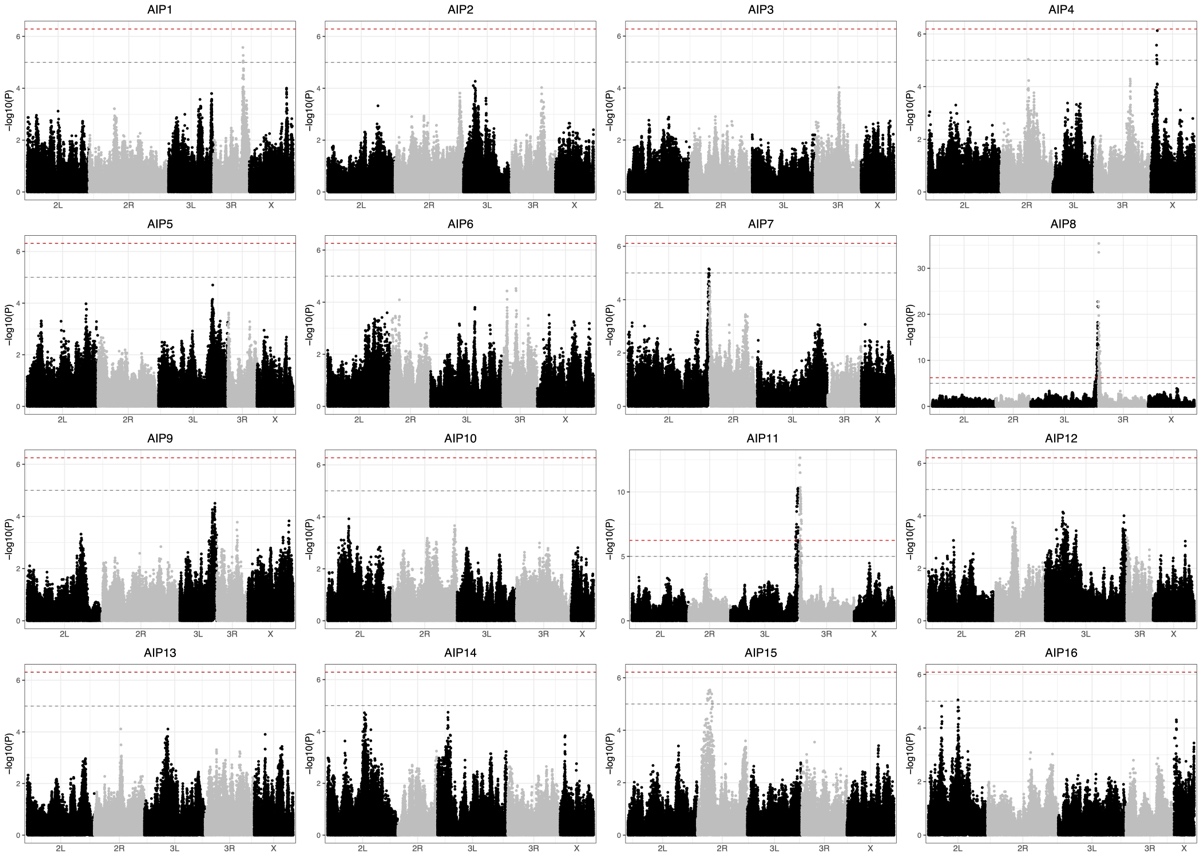

（B）LSD

#### Supplementary Figure 5. Polygenic score analysis for each AIP (cage) under (A) HSD and (B) LSD.

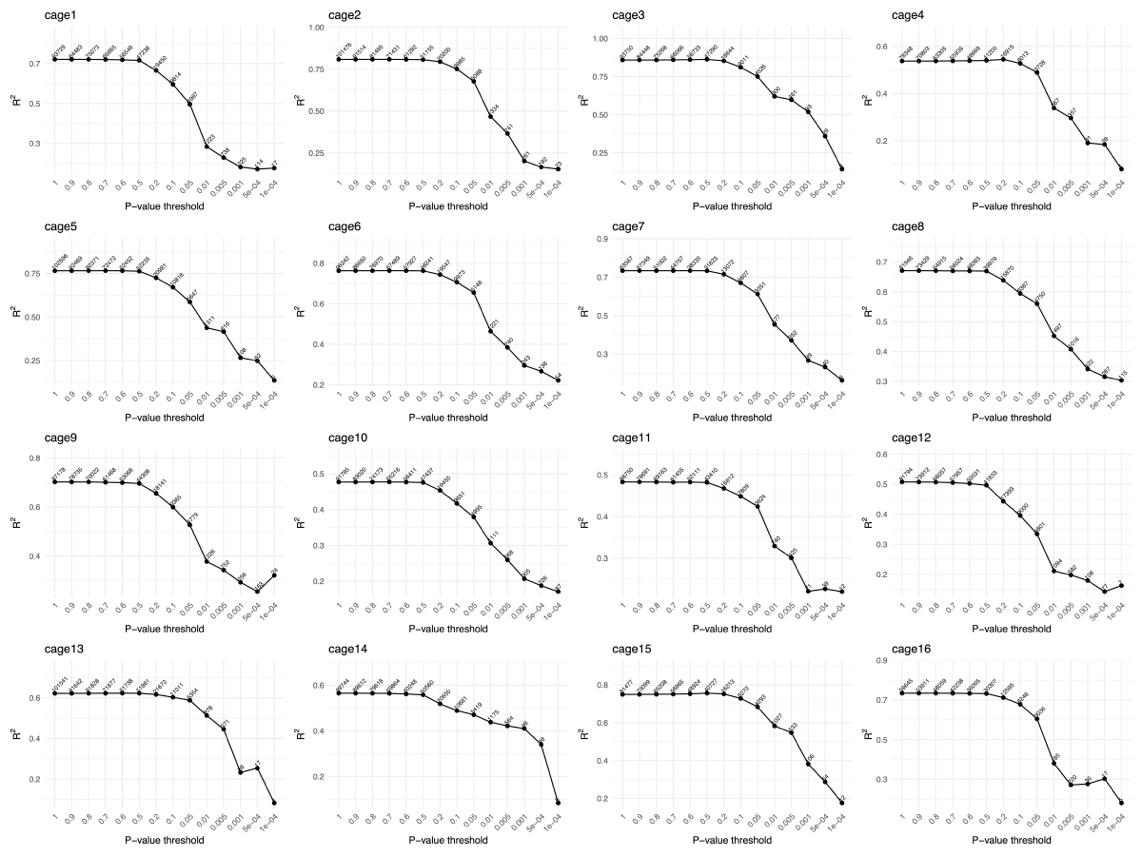

（A）HSD

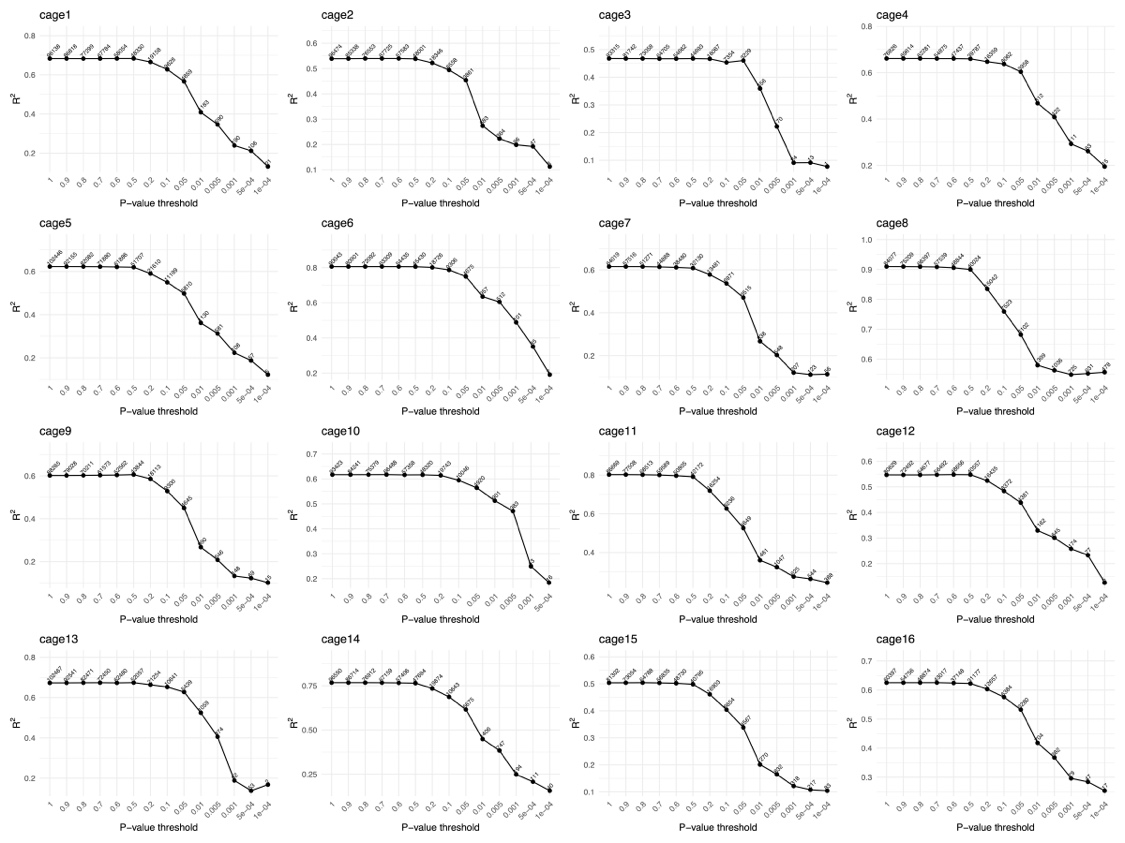

（B）LSD

#### Supplementary Figure 6. Development time distributions between mutant and control groups across diets and sexes.

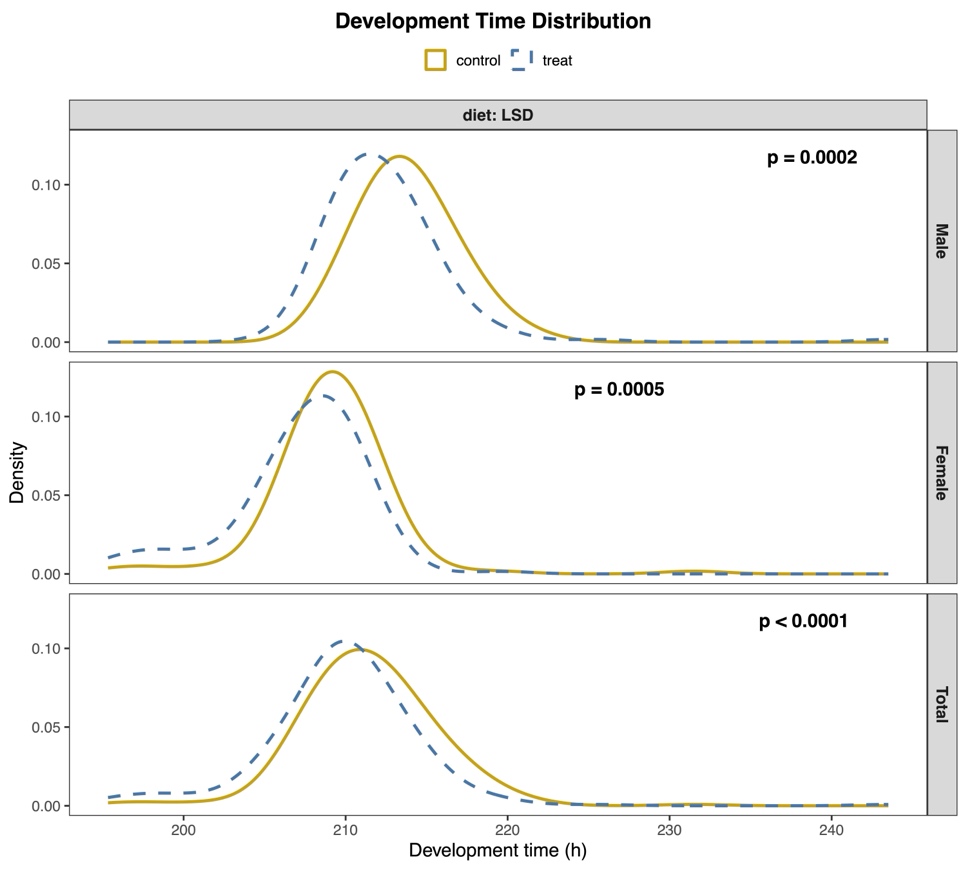

(A) *plx*-RNAi vs control

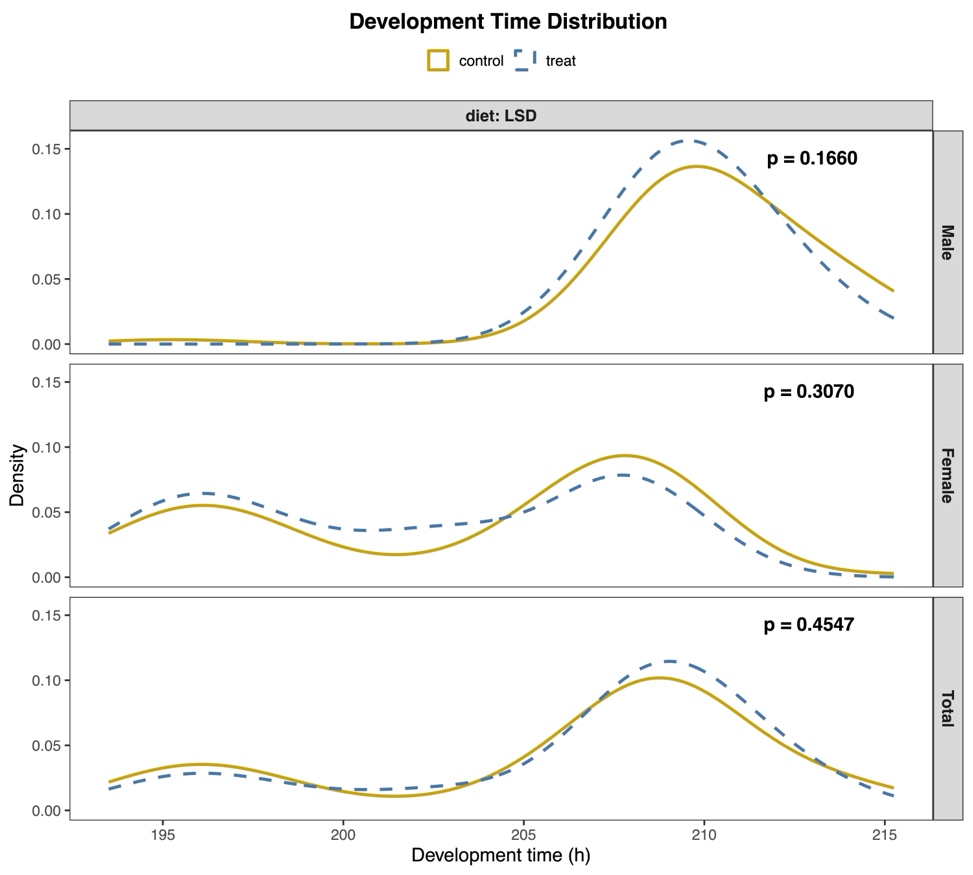

(B) *CG9801*-RNAi vs control

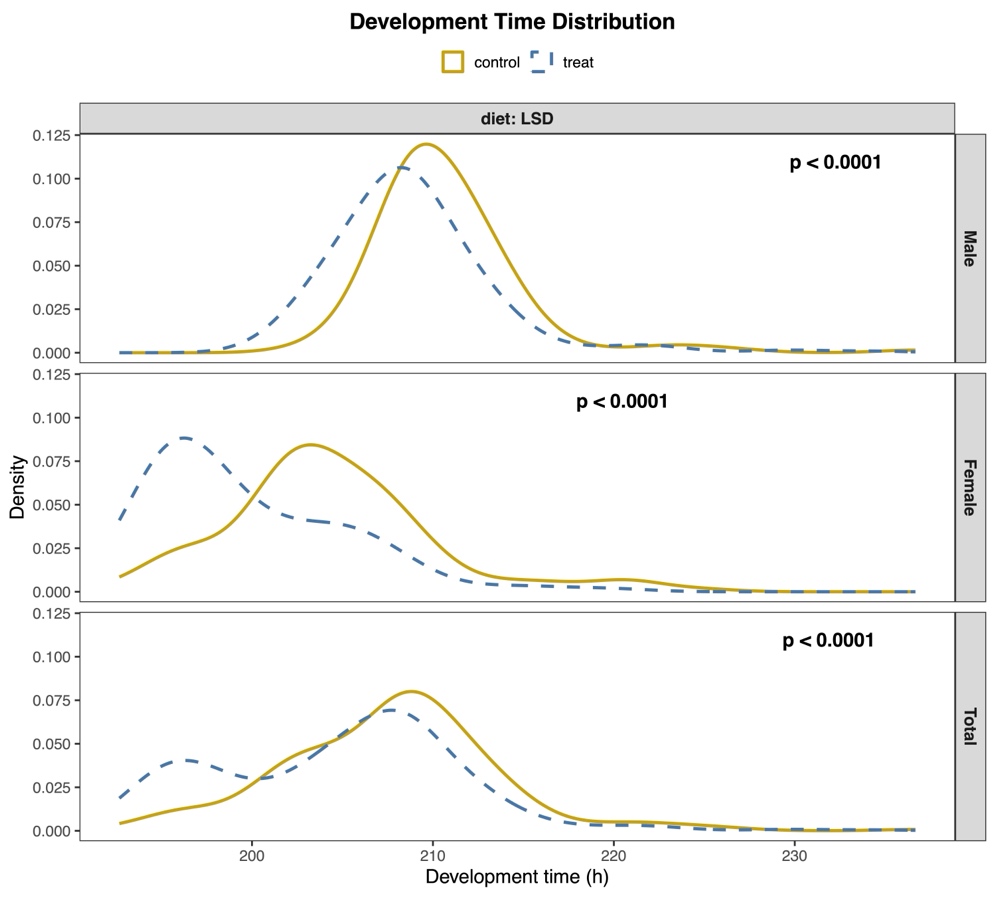

(C) *Pif1B*-RNAi vs control

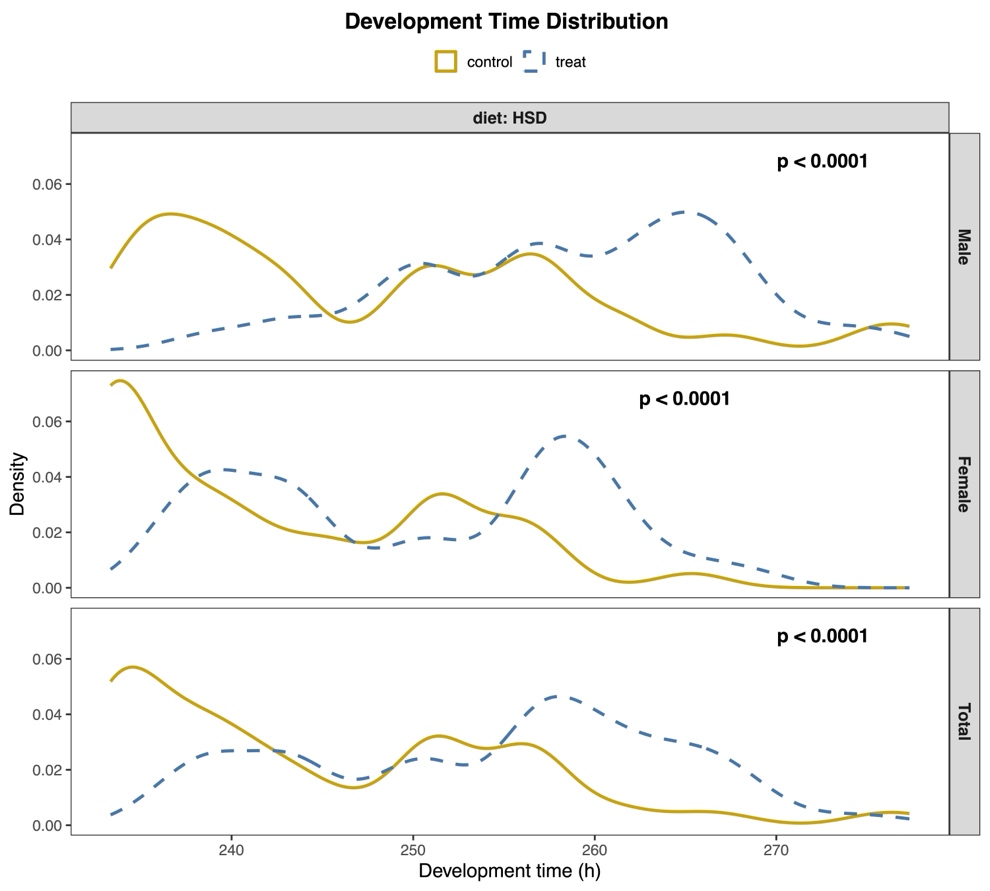

(D) *CG9331*-RNAi vs control

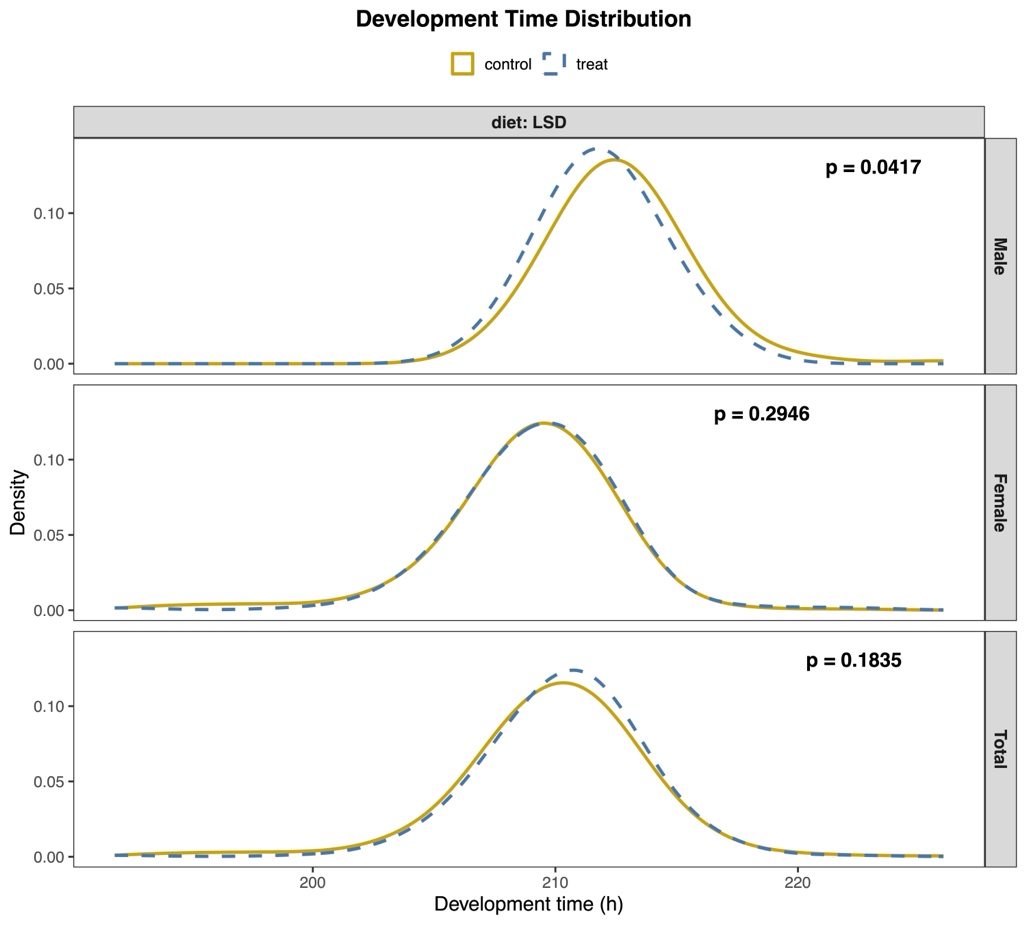

(E) *mtd*-RNAi vs control

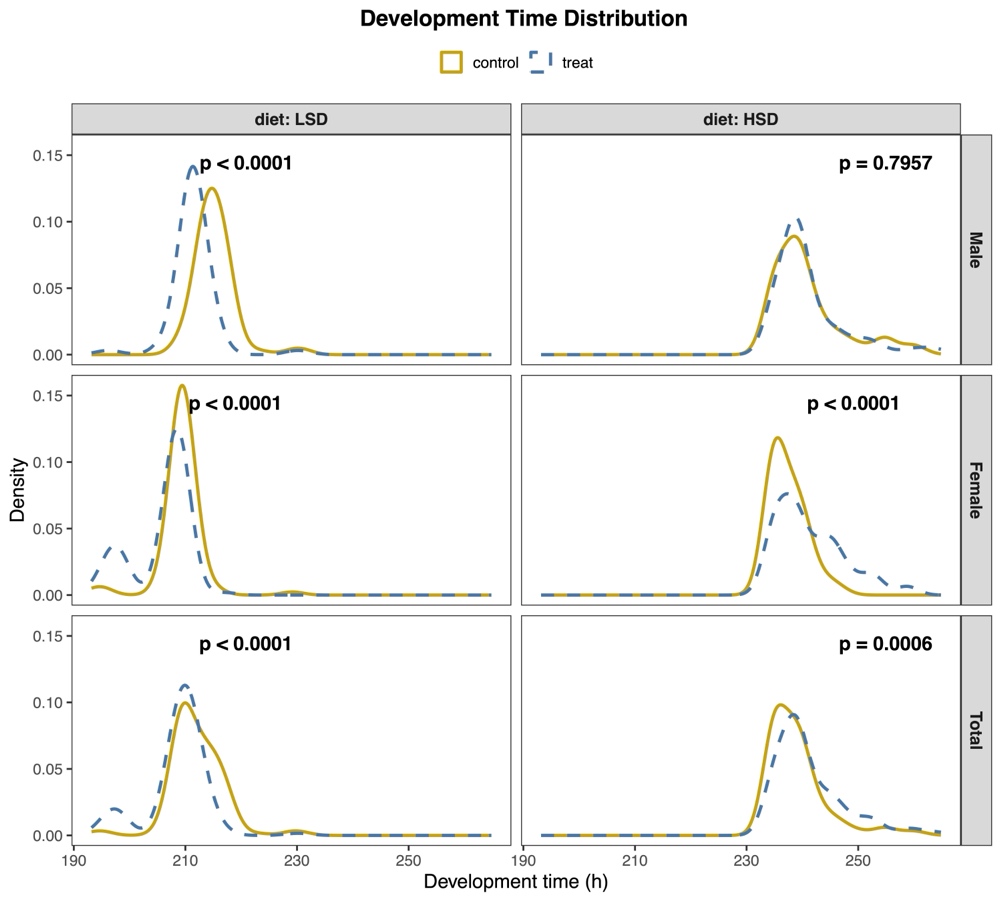

(F) *Cerk*-RNAi vs control

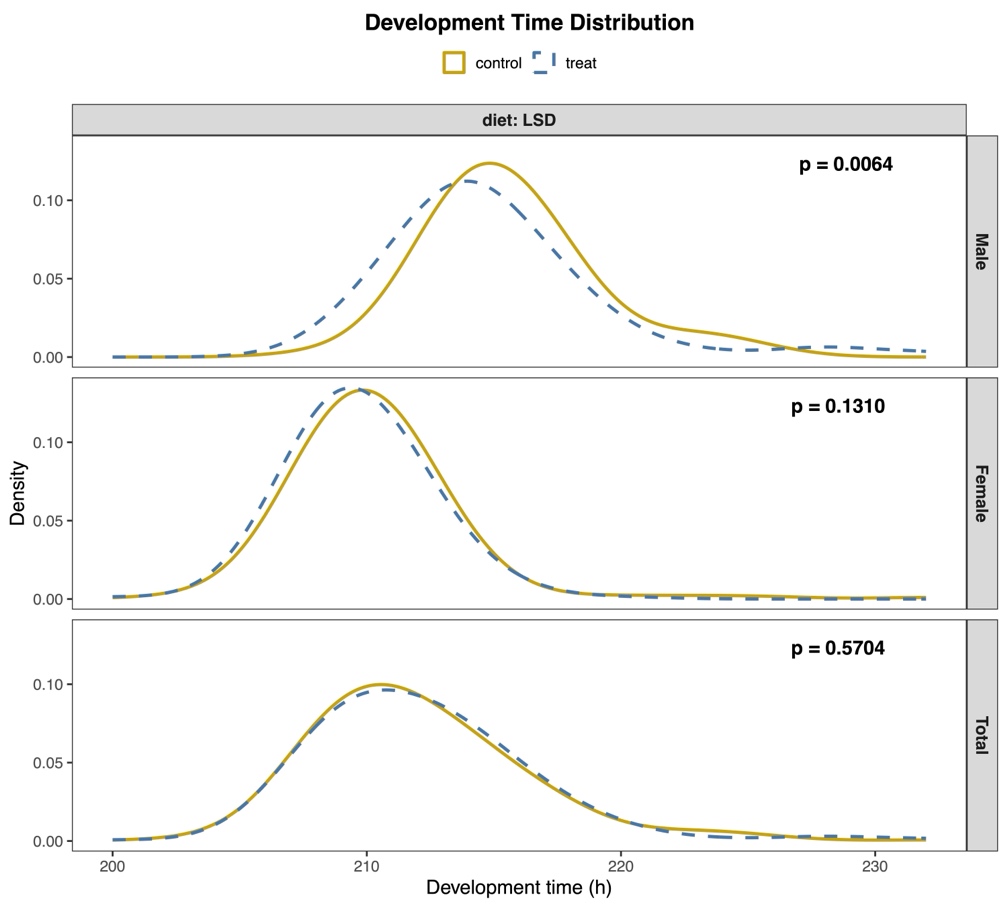

(G) *Sfp79B*-RNAi vs control

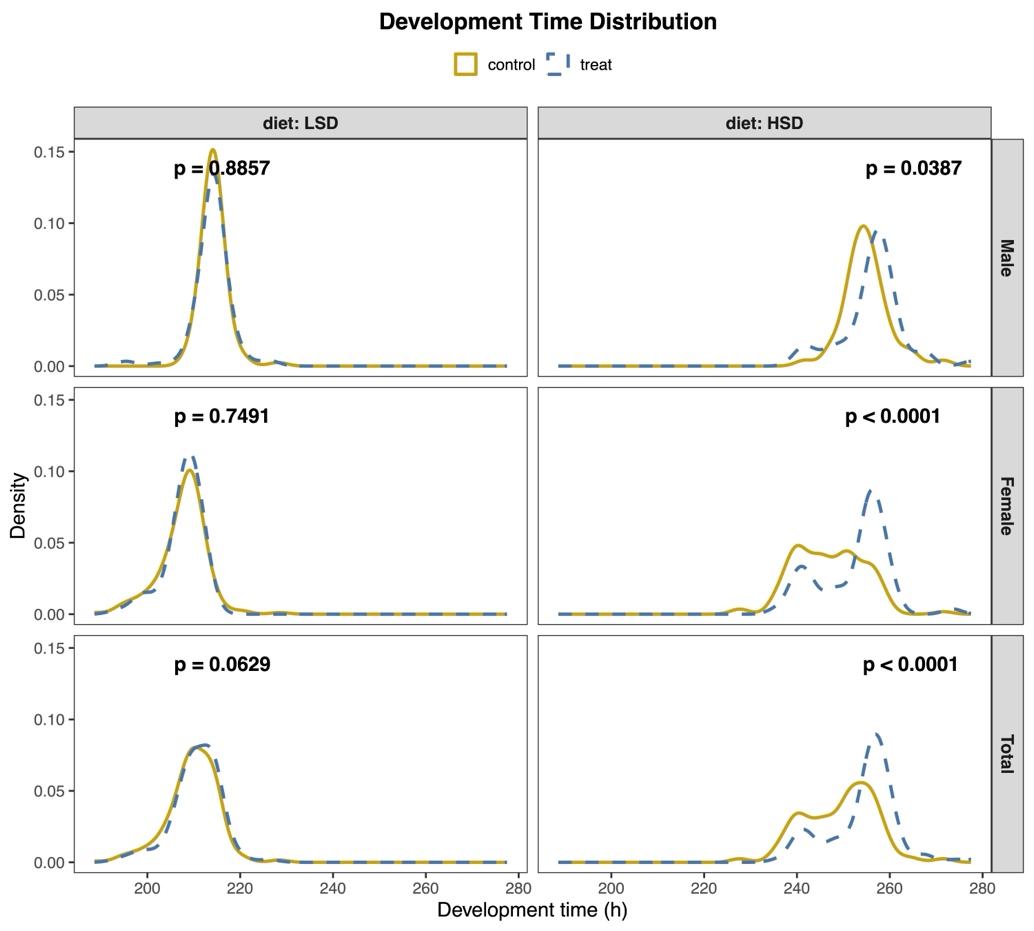

(H) *Pak*-RNAi vs control

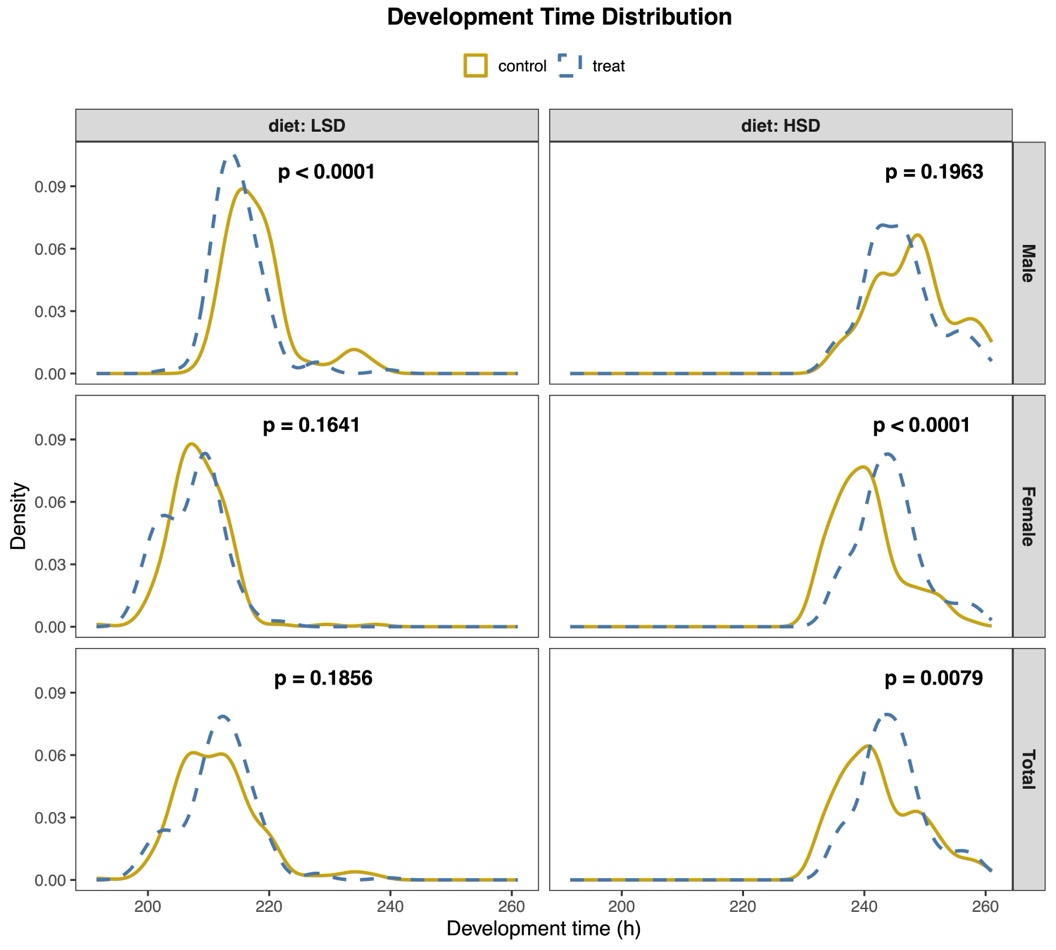

(I) *Eip75B*-RNAi vs control

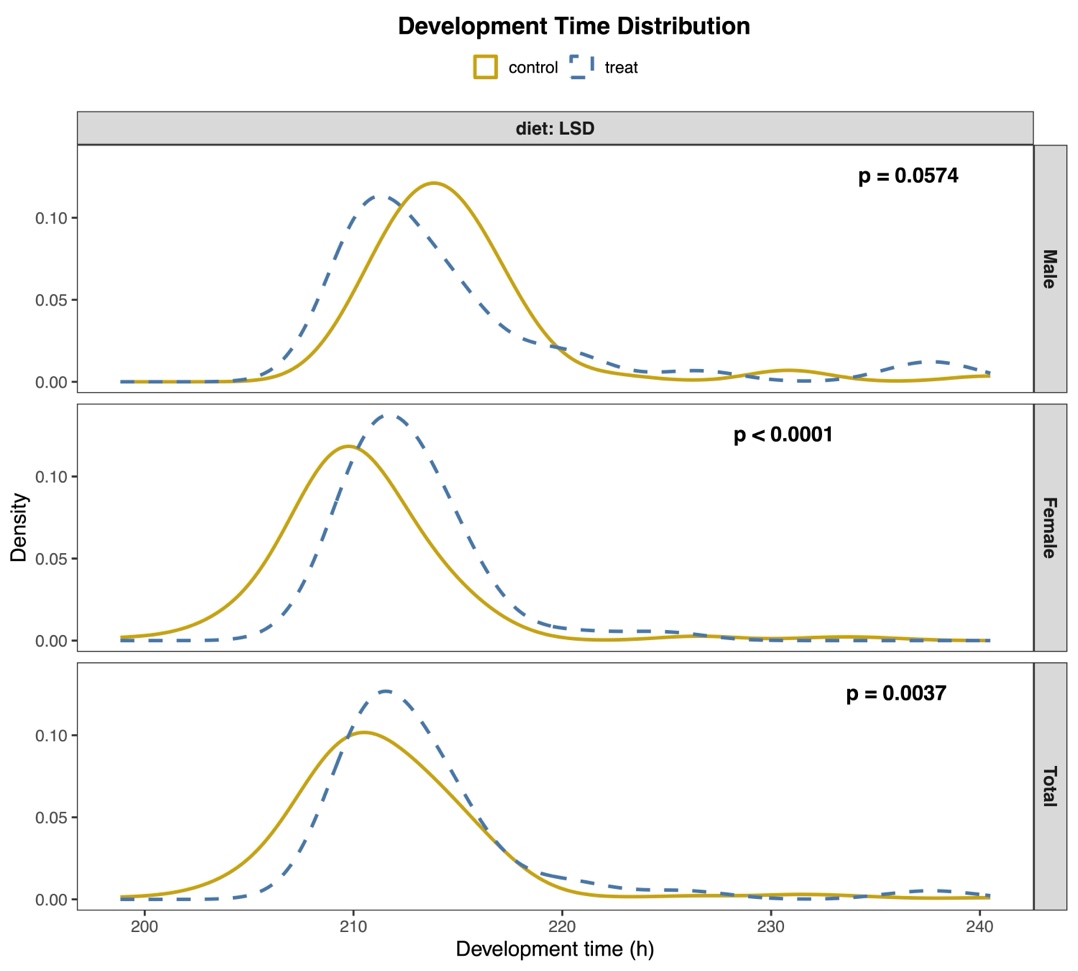

(J) *fz2*-RNAi vs control

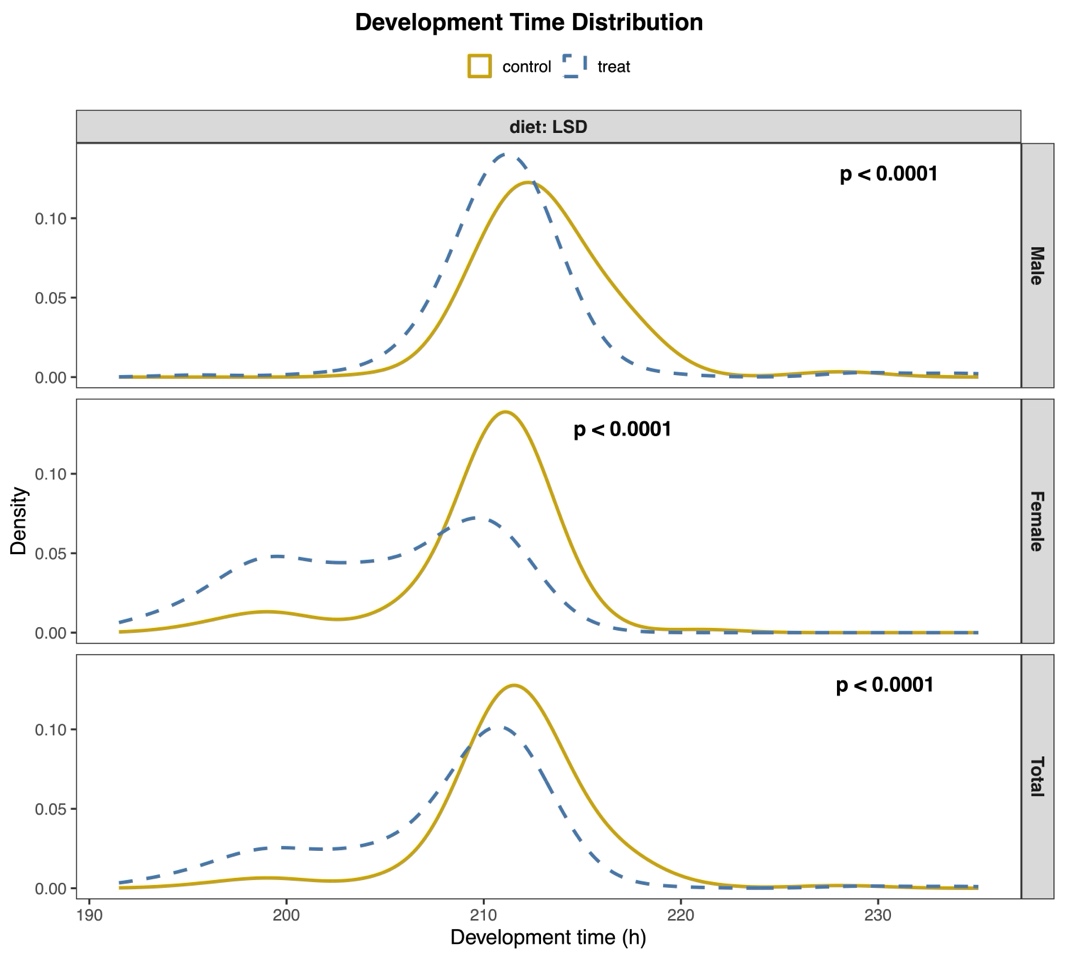

(K) *Dh44*-Overexpression vs control

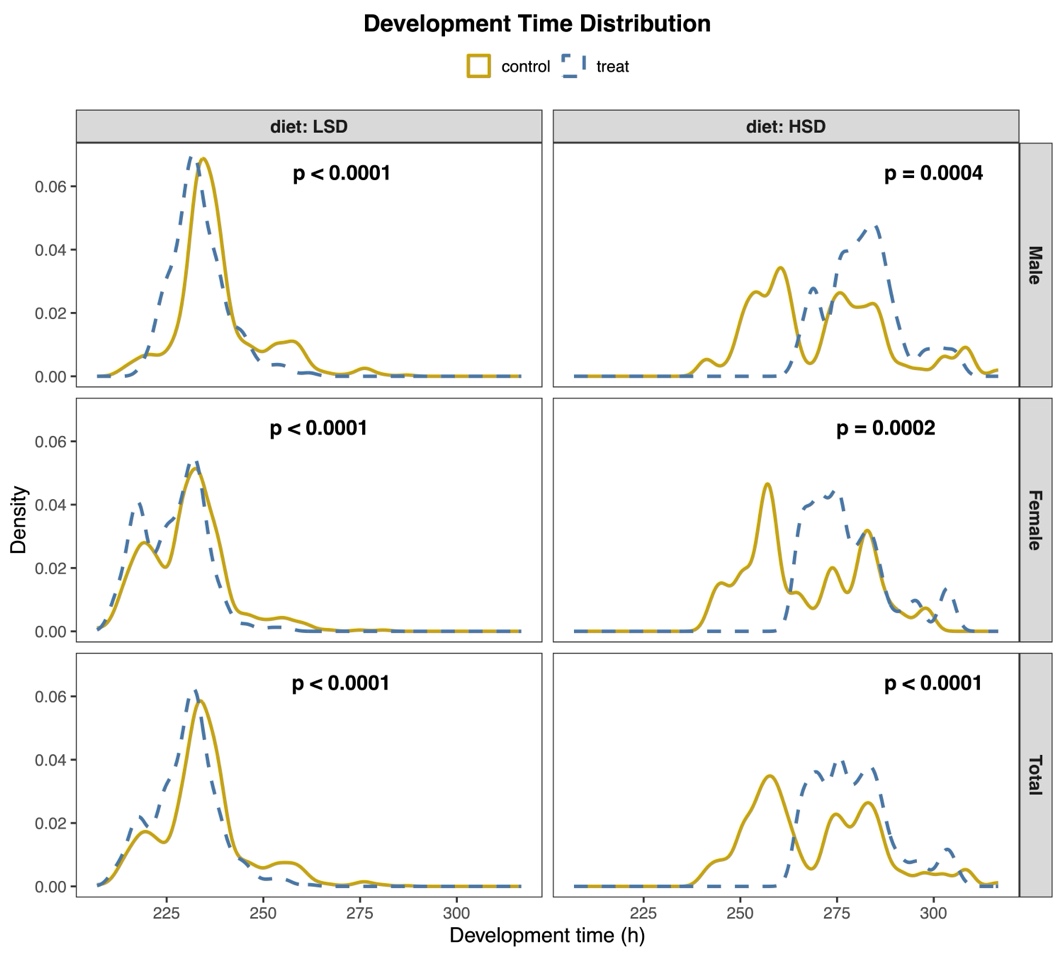

(L) *Tap*-loss-of-function vs control

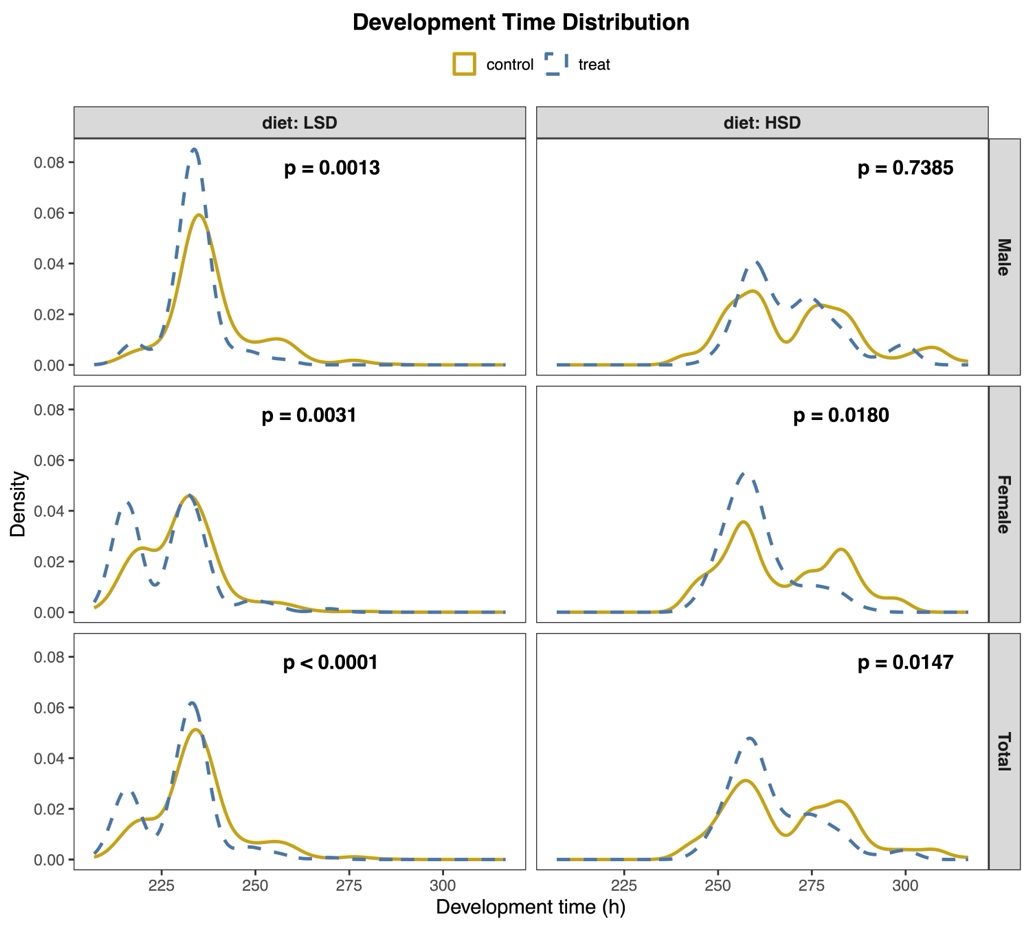

(M) *Cpr76Bd*-loss-of-function vs control

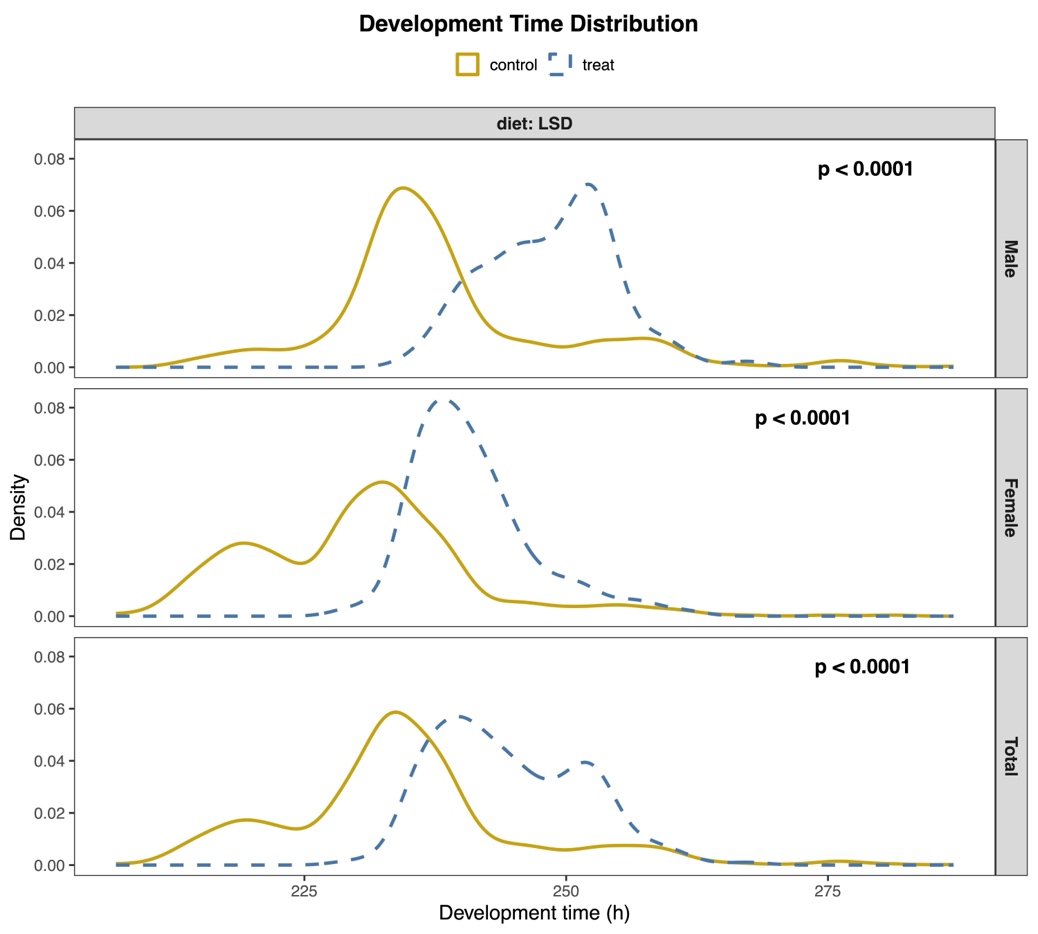

(N) *CG33768*-loss-of-function vs control

#### Supplementary Figure 7. Candidate gene effects on development time across diet and sex.

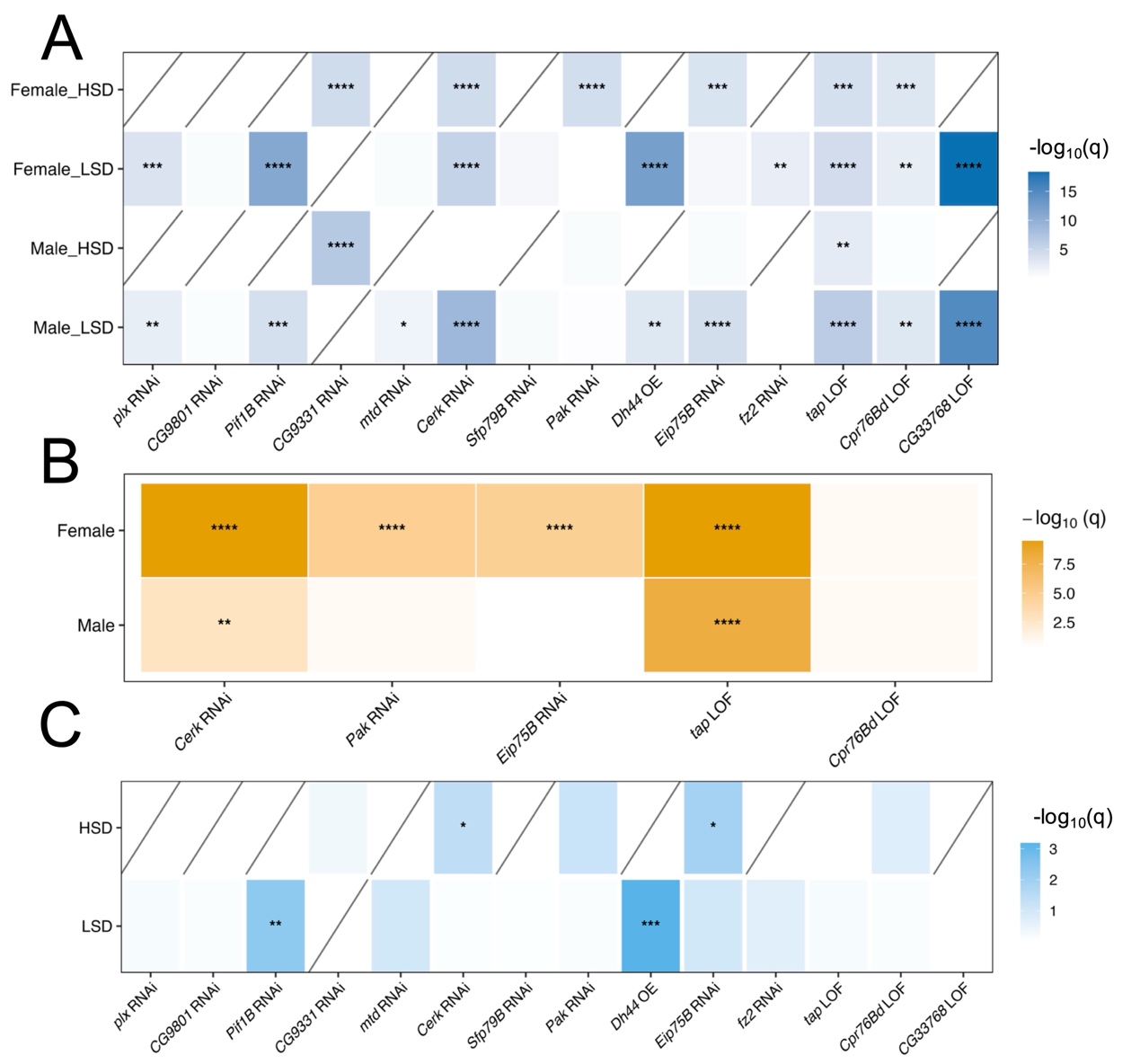

(A) Perturbation effects, testing whether perturbed and control groups differed within each sex-by-diet condition. (B) Diet-specific effects of candidate gene perturbations, testing whether perturbation effects differed between LSD and HSD within each sex. (C) Sex-specific effects of candidate gene perturbations, testing whether perturbation effects differed between males and females within each diet. Color intensity indicates statistical significance, shown as −log10(BH-adjusted q value). Asterisks indicate significance levels after Benjamini–Hochberg correction. Diagonal lines in panels A and C indicate comparisons that were not tested because the required combinations were unavailable. The y axis indicates sex, diet, or sex-by-diet combinations, and the x axis indicates candidate gene perturbation lines. * q < 0.05, ** q < 0.01, *** q < 0.001, **** q < 0.0001.

#### Supplementary Figure 8. Flow chart for the identification of thrifty-like SNPs.

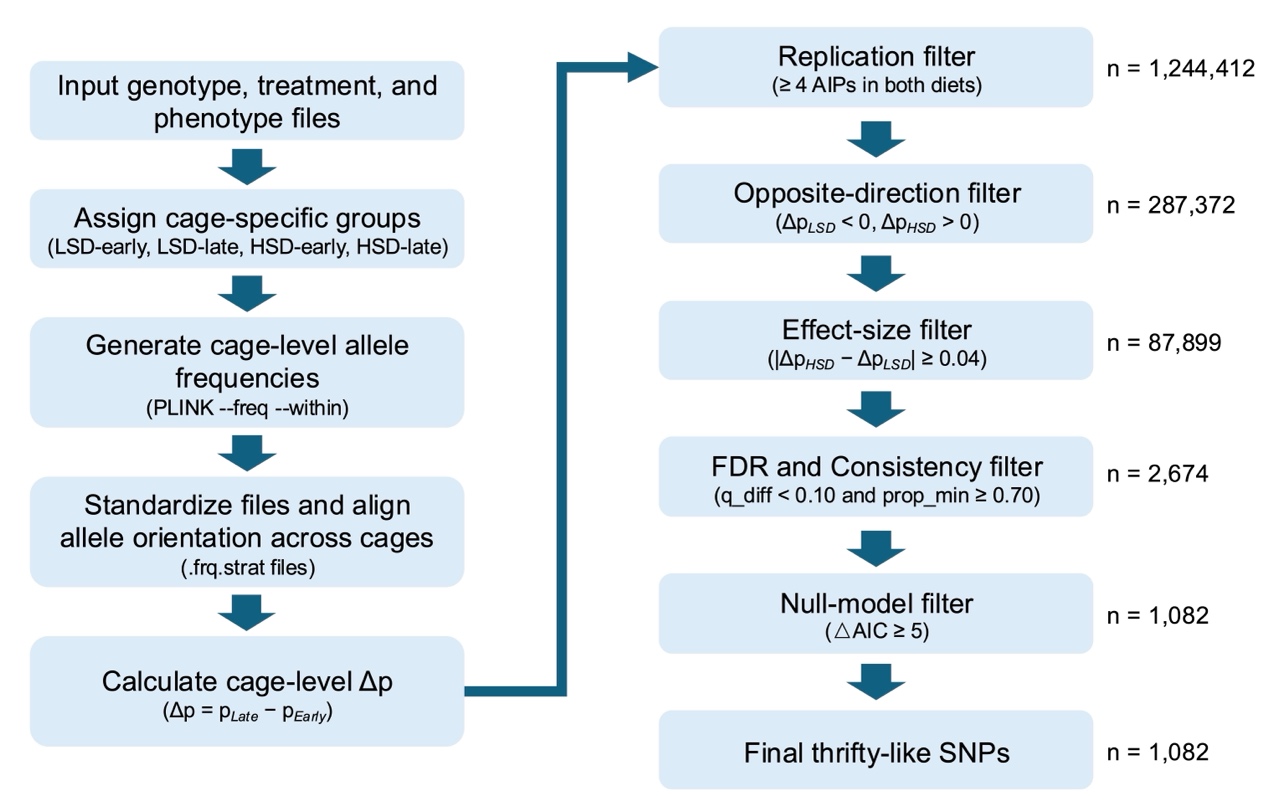

### Supplementary Tables

#### Supplementary Table 1. DGRP lines used for metabolic and life history traits assays (#1-32) and for DRP construction (#1-64).

| DRP Line # | Bloomington Stock # | DGRP Line # |
| --- | --- | --- |
| 1 | 28197 | RAL-440 |
| 2 | 28227 | RAL-761 |
| 3 | 28260 | RAL-897 |
| 4 | 28232 | RAL-790 |
| 5 | 28128 | RAL-45 |
| 6 | 28218 | RAL-703 |
| 7 | 28229 | RAL-776 |
| 8 | 28157 | RAL-228 |
| 9 | 29654 | RAL-320 |
| 10 | 28198 | RAL-441 |
| 11 | 28254 | RAL-879 |
| 12 | 28233 | RAL-796 |
| 13 | 28132 | RAL-75 |
| 14 | 28244 | RAL-822 |
| 15 | 28205 | RAL-508 |
| 16 | 28153 | RAL-195 |
| 17 | 29655 | RAL-321 |
| 18 | 28207 | RAL-531 |
| 19 | 28154 | RAL-217 |
| 20 | 25190 | RAL-380 |
| 21 | 25188 | RAL-375 |
| 22 | 28136 | RAL-91 |
| 23 | 29653 | RAL-229 |
| 24 | 25199 | RAL-639 |
| 25 | 28219 | RAL-716 |
| 26 | 28220 | RAL-721 |
| 27 | 28182 | RAL-370 |
| 28 | 28226 | RAL-757 |
| 29 | 28162 | RAL-256 |
| 30 | 25192 | RAL-399 |
| 31 | 28122 | RAL-21 |
| 32 | 25210 | RAL-859 |
| 33 | 28202 | RAL-491 |
| 34 | 28150 | RAL-177 |
| 35 | 28178 | RAL-356 |
| 36 | 28217 | RAL-646 |
| 37 | 25174 | RAL-208 |
| 38 | 25184 | RAL-357 |
| 39 | 25203 | RAL-732 |
| 40 | 25204 | RAL-765 |
| 41 | 28194 | RAL-392 |
| 42 | 28131 | RAL-73 |
| 43 | 25191 | RAL-391 |
| 44 | 28166 | RAL-309 |
| 45 | 28145 | RAL-149 |
| 46 | 29651 | RAL-40 |
| 47 | 29652 | RAL-57 |
| 48 | 28141 | RAL-129 |
| 49 | 28142 | RAL-136 |
| 50 | 28147 | RAL-158 |
| 51 | 25206 | RAL-786 |
| 52 | 28262 | RAL-907 |
| 53 | 28231 | RAL-787 |
| 54 | 28164 | RAL-280 |
| 55 | 25445 | RAL-365 |
| 56 | 29659 | RAL-513 |
| 57 | 28221 | RAL-727 |
| 58 | 28156 | RAL-227 |
| 59 | 28275 | RAL-235 |
| 60 | 28279 | RAL-887 |
| 61 | 28149 | RAL-176 |
| 62 | 28237 | RAL-805 |
| 63 | 28151 | RAL-181 |
| 64 | 25193 | RAL-427 |

#### Supplementary Table 2. Hidden Markov model (HMM) transition probabilities.

| Founder state at position 1 | Founder state at position 2 | Transition probability |
| --- | --- | --- |
| ii | ii | $\text{[}e^{-r}$ + (1 - $e^{-r}$) $\frac{1}{8}$]^2^ |
|  | jj | $\text{[}$(1 - $e^{-r}$) $\frac{1}{8}$]^2^ |
|  | ij | $\text{[}e^{-r}$ + (1 - $e^{-r}$) $\frac{1}{8}$] $\text{[}$(1 - $e^{-r}$) $\frac{1}{8}$] |
|  | jk | $\text{[}$(1 - $e^{-r}$)$\frac{1}{8}$] $\text{[}$(1 - $e^{-r}$) $\frac{1}{8}$] |
| ij | ij | $\text{[}e^{-r}$ + (1 - $e^{-r}$) $\frac{1}{8}$]^2^ |
|  | ii or jj | $\text{[}e^{-r}$ + (1 - $e^{-r}$)$\frac{1}{8}$] $\text{[}$(1 - $e^{-r}$) $\frac{1}{8}$] |
|  | ik or jk | $\text{[}e^{-r}$ + (1 - $e^{-r}$)$\frac{1}{8}$] $\text{[}$(1 - $e^{-r}$)$\frac{1}{8}$]+ $\text{[}$(1 - $e^{-r}$)$\frac{1}{8}$] $\text{[}$(1 - $e^{-r}$)$\frac{1}{8}$] |
|  | kl | $\text{2 [}$(1 - $e^{-r}$)$\frac{1}{8}$] $\text{[}$(1 - $e^{-r}$)$\frac{1}{8}$] |

#### Supplementary Table 3. ANOVA results for traits measured showing significant line, diet and line by diet interaction effects for most phenotypes.

| Trait | Source | df | MS | F | *P-*value |
| --- | --- | --- | --- | --- | --- |
| Body weight (larvae) | Treatment | 1 | 18.34 | 109.29 | **** |
|  | Fly lines | 27 | 0.26 | 39.18 | **** |
|  | Trt × Fly lines | 27 | 0.17 | 25.27 | **** |
| Body weight (adult) | Treatment | 1 | 2.05 | 63.10 | **** |
|  | Fly lines | 31 | 0.14 | 42.11 | **** |
|  | Trt × Fly lines | 31 | 0.03 | 9.99 | **** |
| Glucose (larvae) | Treatment | 1 | 75.78 | 3.59 | n.s. |
|  | Fly lines | 27 | 50.13 | 20.47 | **** |
|  | Trt × Fly lines | 27 | 21.12 | 8.63 | **** |
| Glucose (adult) | Treatment | 1 | 714.46 | 49.51 | **** |
|  | Fly lines | 31 | 22.82 | 16.00 | **** |
|  | Trt × Fly lines | 31 | 14.43 | 10.12 | **** |
| Glycogen (larvae) | Treatment | 1 | 807.53 | 18.97 | *** |
|  | Fly lines | 27 | 89.52 | 32.21 | **** |
|  | Trt × Fly lines | 27 | 42.58 | 15.32 | **** |
| Glycogen (adult) | Treatment | 1 | 1860 | 34.42 | **** |
|  | Fly lines | 31 | 366.53 | 63.64 | **** |
|  | Trt × Fly lines | 31 | 54.04 | 9.38 | **** |
| Triglyceride (larvae) | Treatment | 1 | 194.23 | 1.71 | n.s. |
|  | Fly lines | 26 | 171.43 | 5.91 | **** |
|  | Trt × Fly lines | 26 | 113.43 | 3.91 | **** |
| Triglyceride (adult) | Treatment | 1 | 483.08 | 1.75 | n.s. |
|  | Fly lines | 23 | 668.53 | 31.95 | **** |
|  | Trt × Fly lines | 23 | 276.54 | 13.22 | **** |
| Survivability | Treatment | 1 | 6.87 | 59.91 | **** |
|  | Fly lines | 28 | 0.25 | 43.04 | **** |
|  | Trt × Fly lines | 28 | 0.11 | 19.56 | **** |
| Longevity | Treatment | 1 | 2008 | 5.46 | * |
|  | Fly lines | 31 | 642.16 | 6.28 | **** |
|  | Trt × Fly lines | 31 | 368.11 | 3.60 | **** |
| Development time | Treatment | 1 | 874689 | 642.11 | **** |
|  | Fly lines | 28 | 2948 | 38.17 | **** |
|  | Trt × Fly lines | 28 | 1362 | 17.64 | **** |

Significant values indicated as: not significant (n.s.), *P <* 0.05 *, *P <* 0.01 **, *P <* 0.001***, *P <* 0.0001****

#### Supplementary Table 4. Between-line (genetic) and within-line (residual) variation, broad sense heritability, genotypic and residual coefficients of variation for phenotypic traits on HSD and LSD.

| Traits | Treatment | Between-line  Variance (Vg) | Within-line  Variance (V_r_) | Total Phenotypic Variance (V_p_) | Heritability (H^2^) | Genotypic Coefficient of Variance (CV_g_) (%) | Residual Coefficient of Variance (CV_r_) (%) |
| --- | --- | --- | --- | --- | --- | --- | --- |
| Body weight (larvae) | LSD | 0.02 | 5.0E-03 | 0.02 | 0.79 | 7.71 | 3.98 |
|  | HSD | 0.06 | 8.3E-03 | 0.07 | 0.89 | 19.99 | 7.17 |
| Body weight  (adult) | LSD | 0.01 | 4.6E-03 | 0.02 | 0.76 | 9.30 | 5.21 |
|  | HSD | 0.02 | 1.9E-03 | 0.02 | 0.90 | 11.69 | 3.79 |
| Glucose  (larvae) | LSD | 5.97 | 2.97 | 8.94 | 0.67 | 25.61 | 18.07 |
|  | HSD | 7.30 | 1.93 | 9.22 | 0.79 | 25.53 | 13.11 |
| Glucose  (adult) | LSD | 2.64 | 0.85 | 3.49 | 0.76 | 20.97 | 11.90 |
|  | HSD | 4.24 | 2.00 | 6.24 | 0.68 | 19.18 | 13.18 |
| Glycogen  (larvae) | LSD | 8.11 | 1.60 | 9.71 | 0.83 | 28.71 | 12.77 |
|  | HSD | 17.20 | 3.95 | 21.16 | 0.81 | 63.60 | 30.49 |
| Glycogen  (adult) | LSD | 24.46 | 3.05 | 27.52 | 0.89 | 50.13 | 17.71 |
|  | HSD | 57.35 | 8.47 | 65.81 | 0.87 | 51.55 | 19.81 |
| Triglyceride  (larvae) | LSD | 25.35 | 13.34 | 38.69 | 0.66 | 48.88 | 35.46 |
|  | HSD | 20.01 | 44.71 | 64.72 | 0.31 | 37.29 | 55.74 |
| Triglyceride  (adult) | LSD | 96.33 | 18.23 | 114.56 | 0.84 | 47.98 | 20.87 |
|  | HSD | 84.31 | 23.61 | 107.93 | 0.78 | 39.42 | 20.86 |
| Survivability | LSD | 5.6E-03 | 4.8E-03 | 0.01 | 0.54 | 8.69 | 8.09 |
|  | HSD | 0.07 | 6.9E-03 | 0.07 | 0.90 | 46.37 | 15.04 |
| Longevity | LSD | 32.35 | 102.68 | 135.03 | 0.24 | 12.77 | 22.76 |
|  | HSD | 128.80 | 101.82 | 230.63 | 0.56 | 28.72 | 25.53 |
| Development  time | LSD | 164.50 | 19.00 | 183.51 | 0.90 | 9.53 | 3.24 |
|  | HSD | 666.56 | 135.43 | 801.98 | 0.83 | 10.56 | 4.76 |

Between-line Variance (V_G_) + Within-line Variance (V_R_) = Total Phenotypic Variance (V_P_); Broad-sense heritability (H²) = V_G_ / (V_G_ + V_R_). VG was estimated as (MSline - V_R_) / n, where n = 5 biological replicates were assumed from SEM = SD / sqrt(n). CV_G_ = 100 × sqrt(V_G_) / X; CV_R_ = 100 × sqrt(V_R_) / X.

**Supplementary Table 5. Statistical summary of perturbation, diet-specific and sex-specific effects on development time.**

| Category | Gene | Perturbation  type | Perturbation effect | Diet-specific effect | Sex-specific effect |
| --- | --- | --- | --- | --- | --- |
| G×E | *tap* | LOF | **** | Male (****)  Female (****) | HSD (n.s.)  LSD (n.s.) |
|  | *Cpr76Bd* | LOF | *** | Male (n.s.)  Female (n.s.) | HSD (n.s.)  LSD (n.s.) |
|  | *Pak* | RNAi | **** | Male (n.s.)  Female (****) | HSD (n.s.)  LSD (n.s.) |
|  | *Eip75B* | RNAi | **** | Male (n.s.)  Female (****) | HSD (*)  LSD (n.s.) |
|  | *Cerk* | RNAi | **** | Male (**)  Female (****) | HSD (*)  LSD (n.s.) |
|  | *CG9331* | RNAi | **** | Male (n.s.)  Female (n.s.) | HSD (n.s.)  LSD (n.s.) |
| HSD | *plx* | RNAi | *** | Male (n.s.)  Female (n.s.) | n.s. |
| LSD | *CG9801* | RNAi | n.s. | Male (n.s.)  Female (n.s.) | n.s. |
|  | *Pif1B* | RNAi | **** | Male (n.s.)  Female (n.s.) | ** |
|  | *mtd* | RNAi | * | Male (n.s.)  Female (n.s.) | n.s. |
|  | *Sfp79B* | RNAi | n.s. | Male (n.s.)  Female (n.s.) | n.s. |
|  | *fz2* | RNAi | ** | Male (n.s.)  Female (n.s.) | n.s. |
|  | *Dh44* | OE | **** | Male (n.s.)  Female (n.s.) | *** |
|  | *CG33768* | LOF | **** | Male (n.s.)  Female (n.s.) | n.s. |

Significant values indicated as: not significant (n.s.), *q <* 0.05 *, *q <* 0.01 **, *q <* 0.001***, *q <* 0.0001****
